# The uptake of metallic nanoparticles in breast cancer cell lines is modulated by the hyaluronan-CD44 axis

**DOI:** 10.1101/2025.02.12.637873

**Authors:** Marie Hullo, Cécile Mathé, Nuria Fonknechten, Céline Lacrouts, Guillaume Piton, Johanna Noireaux, Sylvie Chevillard, Anna Campalans, Emmanuelle Bourneuf

**Author notes:** Corresponding Author Emmanuelle Bourneuf. CEA Fontenay-Aux-Roses. Co-first authors (Marie Hullo, Cécile Mathé).

## Abstract

Radiation enhancement is a promising anti-cancer approach based on a local radiation dose increase due to the presence of metallic nanoparticles (NPs) within cancer cells. Depending on their composition, size and cellular properties, NPs can follow multiple cellular pathways and entry routes. We observed that gold, platinum and TiO_2_ NPs are internalized at higher levels in mesenchymal cells compared to epithelial cells in breast cancer models. A global survey of gene expression between epithelial and mesenchymal cells exposed to 4 different NP types revealed an involvement of membrane structure, and further experiments confirmed that the hyaluronic acid (HA) and its receptor CD44 are mediators of metallic NP uptake into cells. We extended our results to a larger panel of breast cancer cell lines and again showed a preferential uptake of all NPs tested in mesenchymal cells and relying on the HA/CD44 axis. These data provide considerations for the design of NP-based therapies targeting mesenchymal cancer cells, which are often resistant to treatment and correlate with poor prognosis and tumor recurrence.

## INTRODUCTION

Nanomaterials, *i.e.* elements which size ranges from a few to 100nm, represent promising tools for clinical applications. In cancer research, nanoparticles (NPs) are being developed as drug carriers targeting specific cell types, or as sensitizers for radiotherapy (RT). The unique physicochemical properties of nano-sized objects can be harnessed to create multimodal theranostic nanoparticles, that enable for example imaging and radiosensitization simultaneously. Although being one of the pillars of current cancer treatment, radiotherapy can induce severe secondary effects when surrounding healthy tissues receive excessive ionizing radiation. In addition, cancer cells radioresistance, whether intrinsic or acquired during treatment, also hinders the successful elimination of all cancer cells^1^. The development of new radiotherapy modalities (beam design, type of particle, for example) as well as the identification of radiation sensitizing or enhancing molecules is crucial to keep an efficient tumor control while minimizing damage to healthy surrounding tissues, and therefore improve the global therapeutic index of RT^2^. Nanoparticle-enhanced radiotherapy involves using high-Z metallic nanoparticles, which presence in cells will increase the probability of interaction between incident photons and matter^3^. In addition to direct effects of radiations on structures, the massive release of secondary electrons will enhance radiolysis of water molecules, ROS production and subsequent damages to biological molecules, like DNA, lipids and proteins^4^. Ultimately, damage to organelles and DNA will lead to cell death. In addition, in vivo experiments on mouse tumor models injected with hafnium-based NBTXR3 nanoparticles successfully demonstrated the activation of an immune response against cancer cells, suggesting a systemic effect of local radiation dose enhancement^5^. Overall, this approach consisting on a local dose escalation could improve the therapeutic ratio by allowing a global dose de-escalation and healthy tissue sparing, while maintaining an efficient tumor cell killing.

The first *in vivo* proof of concept of nanoparticle-based enhanced radiotherapy was established by Hainfeld in 2004, in an animal model of mammary carcinoma^6^. Since this seminal study, various nanoparticle designs with different metal compositions and surface modifications have been used as radiosensitizers in mouse models^7,8^, and some have even progressed to clinical trials.

An optimal radiation enhancement ratio can be achieved with a maximal uptake of the metallic nanoparticles within cells^9^. This uptake can occur passively, or actively by endocytosis. Endocytosis includes phagocytosis, clathrin and caveolae-mediated endocytosis, and other pathways^10^. Of course, the NP uptake is highly dependent on NPs physico-chemical features, such as size, shape, charge and corona composition^11^. Regarding NP size, permeation will be favored for very small NPs (under 5 nm), while larger NPs require active endocytosis. Therefore, NPs are often functionalized with ligands or other molecules that can target membrane-bound receptors and lead to a specific cellular uptake. Several very comprehensive reviews have discussed NP uptake via different endocytic pathways, highlighting methodological approaches and potential pitfalls that often lead to misinterpretation^9–11^.

Studies specifically focusing on metallic NPs larger than 10 nm have hypothesized different modes of uptake, generally by classical endocytosis. However, cell type variability and NP-cell interactions are often overlooked. Yet tumors represent heterogeneous entities, composed of a complex interplay of many cell types, such as cancer, immune, stromal cells, etc. While cellular heterogeneity offers new treatment opportunities by targeting immune populations for example, the plasticity and dynamic adaptation of cancer cells is critical. For example, when stimulated by growth factors or chemotherapeutic drugs, cells can undergo profound phenotypic and genetic changes, in a process called epithelial-to-mesenchymal transition (EMT). EMT fosters extravasation and metastatic invasion, as well as therapeutic resistance^12^. In breast cancer, mesenchymal or partially mesenchymal (hybrid) cells, albeit in small numbers, are considered to be the most resistant and invasive and are thought to be responsible for a substantial amount of tumor relapses^13^.

To better understand the uptake of nanoparticles in cancer cells, we interrogated various metallic NPs entry in a panel of breast cancer cell lines representing epithelial and mesenchymal phenotypes. A transcriptomic survey allowed to refine the molecular mechanisms explaining that mesenchymal cells take up metallic NPs more effectively, with the involvement of hyaluronan and CD44 pathways. Finally, the findings were confirmed in a broader panel of cell lines, including models of breast cancer EMT, highlighting that the HA/CD44 pathway is hijacked by metallic NPs and that this observation could be exploited for therapeutic purposes.

## RESULTS

### Metallic nanoparticles are differentially internalized across breast cancer cell lines

Numerous cell models are available for breast cancer studies, among which T47D and MDA-MB-231 are commonly used to compare features that could differ between epithelial and mesenchymal like cells. Indeed, T47D cells display a polarized epithelial like phenotype, with a cuboid shape, markers of cell-cell interaction and adhesion, such as E-cadherin (E-cad) and desmosomes and a CD24+/CD44−/E-cad+ molecular phenotype (**Figure 1A**). In contrast, MDA-MB-231 cells are spindle-shaped, highly motile and express genes involved in stem cell features (ZEB1, VIM, α5β1 integrin…), as well as molecular markers CD24−/CD44+/E-cad-. Interestingly, those cells also show a differential resistance to drugs and radiation. Epithelial cells are sensitive to treatments and die mostly by apoptosis, while mesenchymal cells undergo cell cycle arrest but eventually start proliferating again. This study explores the impact of these features on radiation enhancement approaches, starting with their ability to uptake NPs within cells. Taking advantage of physicochemical properties of metallic NPs to observe or quantify them, we investigated their uptake and localization within both T47D and MDA-MB-231 models.

**Figure 1:**
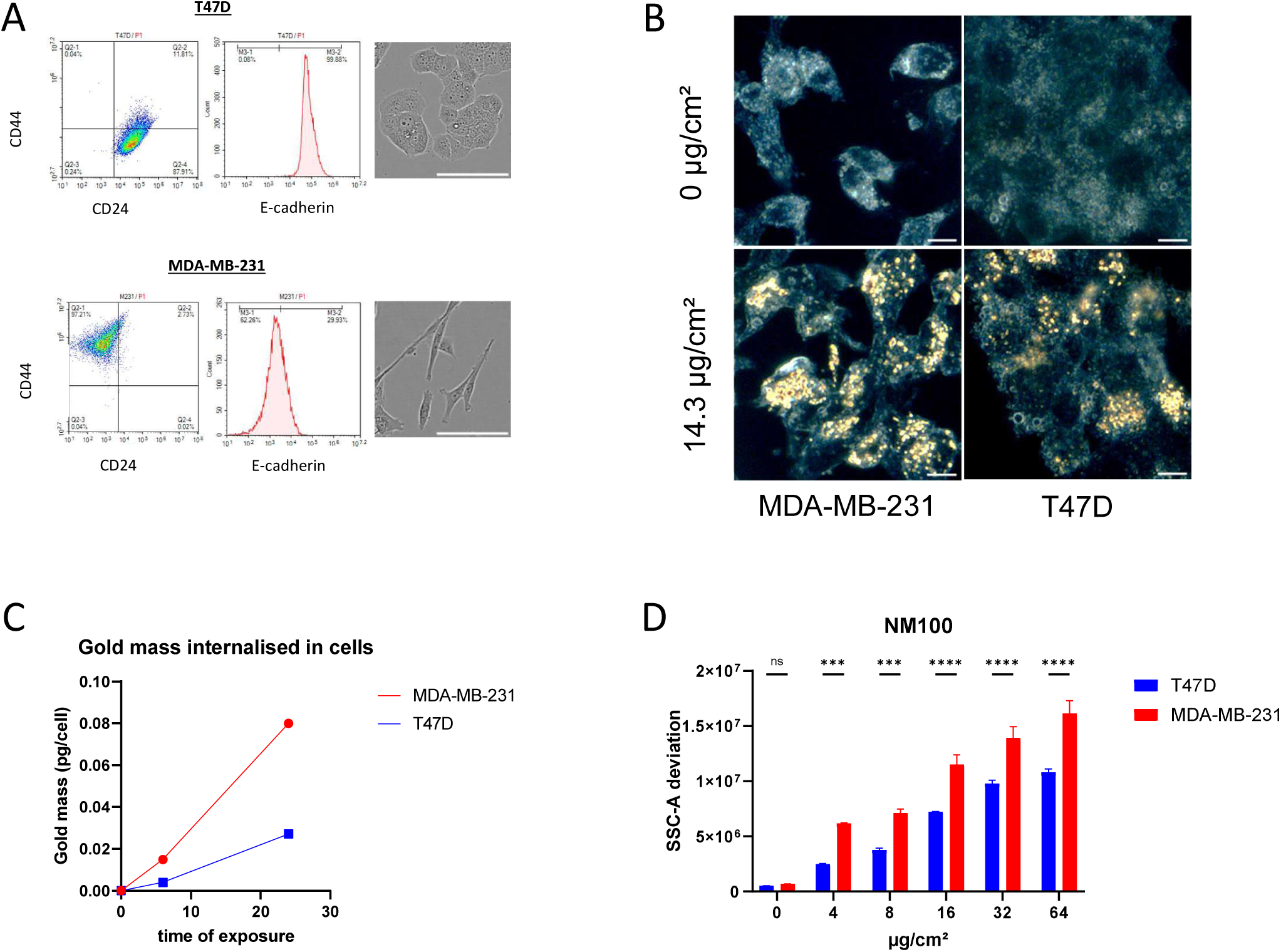
Uptake of metallic nanoparticles by T47D and MDA-MB-231 cells. **A.** Flow cytometry phenotyping of the epithelial (T47D) and the mesenchymal cell lines (MDA-MB-231). Cells were stained with CD44-FITC, CD24-PE and E-cadherin-APC antibodies. The graph represents the mean of data from 10000 cells. The right panels show phase-contrast images of cell lines T47D and MDA-MB-231, scale corresponds to 100 µm. **B.** Darkfield microscopy images of AuNP-NH_2_ (32 nm) internalized in epithelial (T47D) and mesenchymal cells (MDA-MB-231), following a 72 h exposure at 14.3 µg/cm^2^. Scale corresponds to 10 µm. **C.** ICP-MS measure of cytoplasmic gold mass after 6 h and 24 h exposure with 32 µg/cm^2^ AuNPs-NH2 (32 nm), in T47D and MDA-MB-231 cells. **D.** SSC-A deviation signals from T47D and MDA-MB-231 cells exposed to NM100 TiO_2_ NPs with different concentrations (µg/cm²). Cells were exposed for 24 h, and suspensions were subjected to flow cytometry analysis. The graph represents the mean of data from 10000 cells, analyzed in duplicates. Errors bars are SDs. The statistical significance was determined by using the 2-way Anova test. Note: ns, non-significant; *** p < 0.0005 and **** p < 0.0001.

The two cell types were first exposed to 32 nm gold nanoparticles (AuNPs –**Table 1**) for 72 h, and the cells were observed by dark-field microscopy. AuNPs were clearly detected at 72 h exposure, in both cell models, with however more nanoparticles in the MDA-MB-231 cells (**Figure 1B**). Unfortunately, this type of detection is not precise enough to quantify and localise the AuNPs within the cells. In order to objectively measure the amount of gold internalized by the cells, the intracellular gold mass was measured by ICP-MS, 24 hours after exposure to 32 µg/cm² of AuNPs (**Figure 1C**). The absolute amount of gold found in MDA-MB-231 cells corresponded to an average of 0.1 pg/cell, while in T47D cells it was about half. Thus, AuNPs are internalized in both cell lines, and their intracellular levels appear to be different. Next, the uptake of other metallic nanoparticles was tested in the same cell lines to exclude an effect only attributable to gold NPs. The amount of intracellular TiO_2_ NPs (NM-100, with a size of 110 nm, **Table 1**) was evaluated by flow cytometry, after 24 h exposure to the NPs and two washes to avoid a potential bias due to membrane-bound NPs. The measure of side scatter signal (SSC) deviation, although not perfectly quantitative, is a convenient proxy for the evaluation of TiO_2_ NPs, AgNPs (silver NPs) and gold NPs cellular uptake^14–16^. The results are shown in **Figure 1D**, where an increase in the SSC deviation can be observed in both cell lines, proportional to the TiO_2_ NPs concentration. Again, a higher uptake was observed in MDA-MB-231 compared to T47D cells. In a previous report we showed an equivalent result for PtNPs (platinum NPs) internalized in MDA-MB-231 and T47D cells^17^. As early as after 2 h of exposure, PtNPs accumulate in structures resembling mutivesicular bodies, providing insight into the subcellular localization of PtNPs. The ICP-MS method further demonstrated a higher mass of platinum in MDA-MB-231 compared to T47D cells. Finally, we already showed a preferential uptake of gold NPs in MDA-MB231 compared to T47D cells, measured by sp-ICPMS^18^.

**Table 1:**
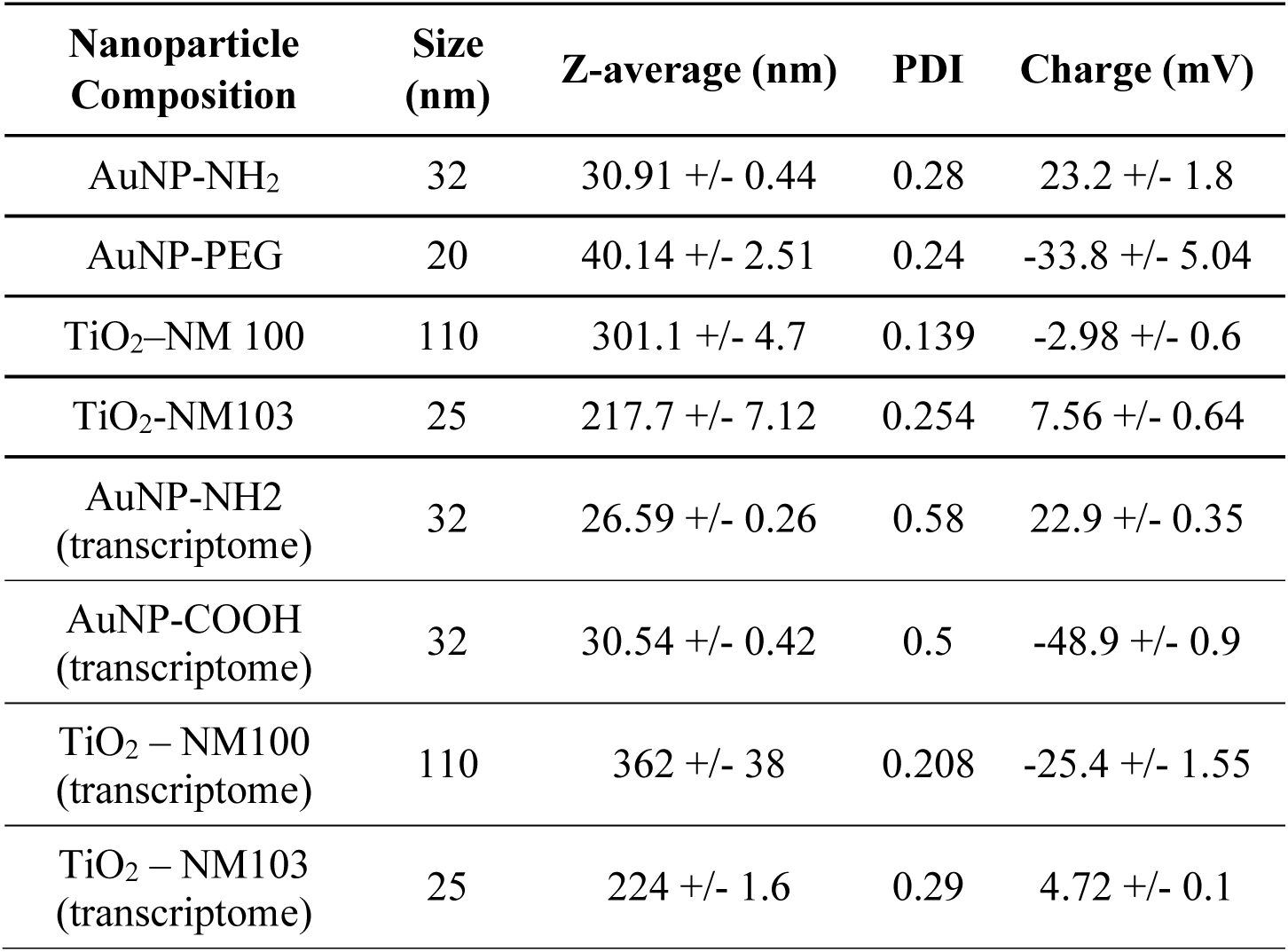
Physico chemical features of the different nanoparticles suspensions used in the study. Measures were performed with a Zetasizer equipment, and analysed with the Malvern Zetasizer software, in triplicates.

Taken together, these results showed a concentration and time-dependent uptake of several types of NPs in two breast cancer cell lines. Furthermore, MDA-MB-231 cells internalized a higher amount of NPs compared to T47D cells, irrespective of the NP.

Since T47D and MDA-MB-231 cells are derived from tumours excised from different patients, the observed difference could be due to variable pathways or metabolic alterations between the cells. Therefore, it seems important to test such an observation in cellular models with the same genetic background. Therefore, we took advantage of two epigenetic models, namely HMLE^19^ and MCF10A cells, which illustrate the breast epithelial-mesenchymal transition *in vitro*. Initially, both cell types exhibit an epithelial-like phenotype with CD24^+^/CD44^-^ labelling, and express high levels of E-cadherin (**Figure 2A**). Upon exposure to chronic doses of EMT inducer, here TGFβ (Transforming Growth Factor – β), HMLE-E and MCF10A-E cells can undergo a reversible epithelial to mesenchymal transition. The resulting mesenchymal cells (HMLE-M and MCF10A-M) show a mesenchymal phenotype, with CD24^-^/CD44^+^ markers and a more fusiform and proliferative phenotype (**Figure 2A**).

**Figure 2:**
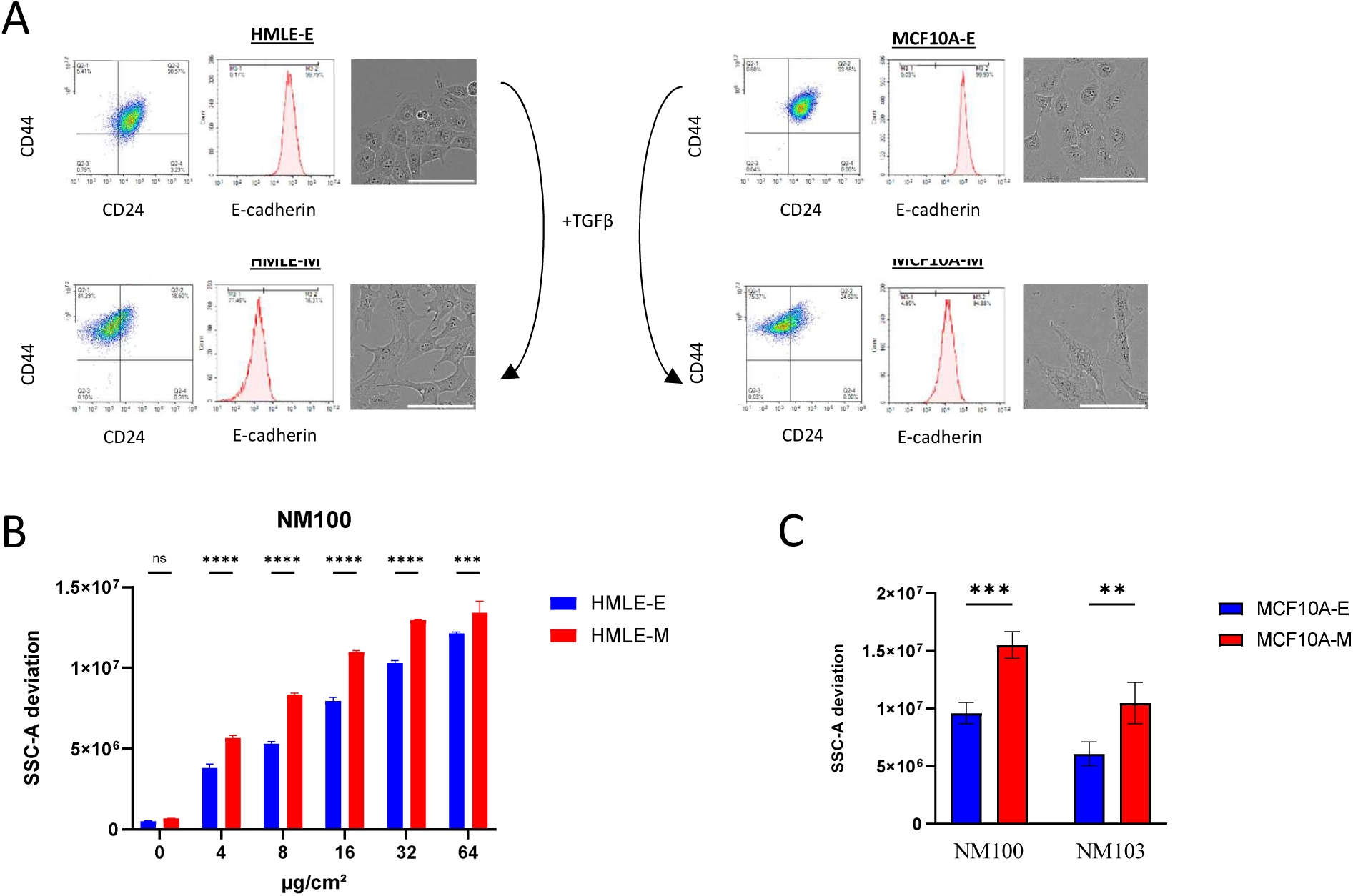
Uptake of TiO_2_ nanoparticles in two breast cancer mesenchymal transition. **A.** Models of induced epithelial to mesenchymal transition *in vitro* with TGFβ. All cell lines were phenotyped by flow cytometry with CD24-PE, CD44-FITC, and E-cadherin-APC antibodies. The right panels show contrast-phase images of the 4 cell lines, magnification x10. Scale bar 100 µm. **B.** SSC-A deviation signals from HMLE-E and M cells exposed to NM100 TiO_2_ NPs with different concentrations (µg/cm²). Cells were exposed for 24 h, and suspensions were subjected to flow cytometry analysis. The graph represents the mean of data from 10000 cells, and in duplicates, error bars are SDs. **C.** SSC-A deviation signals from MCF10A-E and M cells exposed to NM100 and NM103 TiO_2_ NPs with 16 µg/cm². Cells were exposed for 24 h, and suspensions were subjected to flow cytometry analysis. The graph represents the mean of data from 10000 cells, and in triplicates, error bars are SDs. The statistical significance was determined by using the 2-way Anova test and Sidak correction. Note: ns, non-significant; ** p < 0.01 *** p < 0.001 and **** p < 0.0001.

To assess the impact of EMT status in NP uptake, we use TiO_2_ NPs of two different sizes (NM100 - size 110 nm and NM103 - size 25 nm). **Figure 2B** illustrates that the signal deviation in side scatter (SSC-A) increased gradually with the concentration of the NM-100 suspension. In addition, the uptake in HMLE-M was higher than in HMLE-E cells across all concentrations studied. Another experiment investigating the impact of cell confluence on uptake also revealed a higher deviation in HMLE-M than HMLE-E cells. Furthermore, while a higher uptake was evidenced at lower cell densities, the difference between epithelial and mesenchymal remained significant (**Supplementary Figure 1**). Other experiments also showed that the cell surface difference between HMLE-E and M cells was barely significant at high confluence (data not shown). Thus, although cell confluence can have a tremendous impact on cell behaviour, especially in these epigenetic transition models, our results show that metallic NPs uptake is consistently different between epithelial and mesenchymal cells. Regarding MCF10A cells, the SSC-A deviation was 50% higher in MCF10A-M compared to MCF10A-E cells, upon exposure to a fixed concentration of NM100 and NM103 TiO_2_ nanoparticles (**Figure 2C**). These results show that both MCF10A and HMLE epigenetic models internalize more metallic nanoparticles when they are in their mesenchymal state.

The differential uptake of NPs was investigated in cellular models of EMT from other tissues. A549 cells, representing lung epithelial cells, undergo EMT when treated with TGFβ (**Supplementary Figure 2A**). However, the CD44 expression in A549 and A549-M lines as observed by flow cytometry is heterogeneous and does not differ significantly between the lines. In addition, the uptake of TiO_2_ or gold NPs is not statistically different between epithelial and mesenchymal cells (**Supplementary Figure 2B**). Similarly, RWPE1 is a prostatic epithelial cell line, from which WPE1-NB26 mesenchymal cells were derived. The WPE1-NB26 cell line, which exhibits high invasiveness and mesenchymal features, shows no difference in CD44 expression compared to the RWPE1 cell line **(Supplementary Figure 3A**). No significant difference can be observed in NP uptake between epithelial and mesenchymal prostate lines (**Supplementary Figure 3B**), although an increase in NP uptake is likely in the WPE1-NB26 cell line.

To summarize, we show that a differential pattern of internalization exists between breast cancer cells, and that this difference is reproducible even with NPs harbouring different properties, such as a various range of sizes. In addition, in all tested conditions, uptake was more important in mesenchymal cell lines compared to epithelial ones, regardless of confluence or genetic background.

### Transcriptomic analysis of a differential uptake of nanoparticles between epithelial and mesenchymal cells

To explore the pathways involved in the differential internalization of nanoparticles, HMLE epithelial and mesenchymal cells were exposed to several NPs and gene expression profile was analyzed using microarrays. The HMLE model was chosen to avoid potential genetic background interference from the cell lines. Four different types of gold and TiO_2_ NPs were selected with the objective of finding common molecular mechanism to the uptake of NPs of variable size, coating, charge and composition (as mentioned in **Table 1**). HMLE-E and HMLE-M cells were exposed to a NP concentration of 16 µg/cm² for 24 or 48 hours. Differential gene expression was further analyzed taking into account only genes consistently altered across nanoparticles and timepoints, revealing potential candidates explaining a consistent difference in NPs uptake between epithelial and mesenchymal cells. A list of 541 differentially expressed genes was obtained (FC<-2 or >2 and FDR-corrected p-value<0.001; **Supplementary table 1**), and **tables 2** and **3** show the most overexpressed genes in mesenchymal and epithelial cells, respectively. In both cases, the experiment revealed genes that are usually highly expressed in the specific epithelial and mesenchymal breast cancer cells, and found in many other studies. For example, *ZEB1* (FC=57.4, p-val=1.3×10^−13^) and *SERPINE1* (FC=64.57, p-val=5.7×10^−13^), overexpressed in mesenchymal cells, are markers of EMT^20,21^. Conversely, expression of *EPCAM* (FC=1609, p-val=5.06×10^−19^), *ESRP1* (FC=1162, p-val=6.79×10^−16^), and *CDH1* (FC=90, p-val=3.62×10^−14^), is not surprisingly much higher in epithelial cells. These findings align with commonly used markers of an epithelial state, such as E-cadherin ^20^.

**Table 2:** List of the most significantly overexpressed genes in Mesenchymal cells compared to epithelial cells.

| Gene IDs | Gene Name | Description | Fold Change | P-val | FDR P-val |
| --- | --- | --- | --- | --- | --- |
| NM_006329 | FBLN5 | fibulin 5 | 1068.81 | $5.62 \times 10^{-14}$ | $7.09 \times 10^{-11}$ |
| NM_001135934 | POSTN | periostin, osteoblast specific factor | 559.85 | $2.17 \times 10^{-14}$ | $4.65 \times 10^{-11}$ |
| NM_001191322 | GREM1 | gremlin 1, DAN family BMP antagonist | 169.44 | $1.05 \times 10^{-13}$ | $1.11 \times 10^{-10}$ |
| NM_001142474 | NREP | neuronal regeneration related protein | 105.59 | $2.50 \times 10^{-12}$ | $1.62 \times 10^{-09}$ |
| NM_005909 | MAP1B | microtubule associated protein 1B | 67.82 | $2.73 \times 10^{-14}$ | $5.32 \times 10^{-11}$ |
| NM_000602 | SERPINE1 | serpin peptidase inhibitor, clade E, member 1 | 64.57 | $5.72 \times 10^{-13}$ | $4.72 \times 10^{-10}$ |
| NM_001128128 | ZEB1 | zinc finger E-box binding homeobox 1 | 57.4 | $1.34 \times 10^{-13}$ | $1.31 \times 10^{-10}$ |
| NM_001311160 | THY1 | Thy-1 cell surface antigen | 48.57 | $1.59 \times 10^{-12}$ | $1.06 \times 10^{-09}$ |
| NM_001552 | IGFBP4 | insulin like growth factor binding protein 4 | 35.9 | $1.34 \times 10^{-14}$ | $4.10 \times 10^{-11}$ |
| NM_001159 | AOX1 | aldehyde oxidase 1 | 33.26 | $5.20 \times 10^{-12}$ | $3.02 \times 10^{-09}$ |
| NM_001167738 | NAV1 | neuron navigator 1 | 28.77 | $6.24 \times 10^{-12}$ | $3.48 \times 10^{-09}$ |
| NM_014391 | ANKRD1 | ankyrin repeat domain 1 (cardiac muscle) | 28.67 | $1.35 \times 10^{-12}$ | $9.62 \times 10^{-10}$ |
| NM_003713 | PLPP3 | phospholipid phosphatase 3 | 25.03 | $2.80 \times 10^{-11}$ | $1.17 \times 10^{-08}$ |
| NM_001243286 | PVRL3 | poliovirus receptor-related 3 | 24.75 | $1.93 \times 10^{-14}$ | $4.59 \times 10^{-11}$ |
| NM_001311175 | TNFAIP8L3 | tumor necrosis factor, alpha-induced protein 8-like 3 | 22.8 | $6.65 \times 10^{-12}$ | $3.48 \times 10^{-09}$ |
| NM_001290263 | DOCK10 | dedicator of cytokinesis 10 | 16.73 | $2.57 \times 10^{-11}$ | $1.10 \times 10^{-08}$ |
| NM_005328 | HAS2 | hyaluronan synthase 2 | 16.5 | $6.63 \times 10^{-12}$ | $3.48 \times 10^{-09}$ |
| NM_001134999 | FERMT2 | fermitin family member 2 | 7.37 | $9.51 \times 10^{-14}$ | $1.07 \times 10^{-10}$ |
| NM_014729 | TOX | thymocyte selection-associated high mobility group box | 6.39 | $7.25 \times 10^{-12}$ | $3.63 \times 10^{-09}$ |
| NM_004305 | BIN1 | bridging integrator 1 | 4.09 | $1.41 \times 10^{-12}$ | $9.79 \times 10^{-10}$ |

**Table 3:** List of the most significantly overexpressed genes in Epithelial cells compared to mesenchymal cells.

| Public Gene IDs | Gene Name | Description | Fold Change | P-val | FDR P-val |
| --- | --- | --- | --- | --- | --- |
| NM_052886 | MAL2 | mal, T-cell differentiation protein 2 (gene/pseudogene) | 2375.39 | $3.70 \times 10^{-18}$ | $3.97 \times 10^{-14}$ |
| NM_002354 | EPCAM | epithelial cell adhesion molecule | 1609.15 | $5.06 \times 10^{-19}$ | $1.09 \times 10^{-14}$ |
| NM_001034915 | ESRP1 | epithelial splicing regulatory protein 1 | 1162.17 | $6.79 \times 10^{-16}$ | $3.64 \times 10^{-12}$ |
| NM_001077491 | KLK5 | kallikrein related peptidase 5 | 584.71 | $2.62 \times 10^{-13}$ | $2.36 \times 10^{-10}$ |
| NM_000494 | COL17A1;<br>MIR936 | collagen, type XVII, alpha 1; microRNA 936 | 180.31 | $4.37 \times 10^{-14}$ | $6.25 \times 10^{-11}$ |
| NM_001012964 | KLK6 | kallikrein related peptidase 6 | 175.08 | $9.66 \times 10^{-12}$ | $4.71 \times 10^{-09}$ |
| NM_021978 | ST14 | suppression of tumorigenicity 14 (colon carcinoma) | 150.73 | $5.74 \times 10^{-15}$ | $2.05 \times 10^{-11}$ |
| NM_004360 | CDH1 | cadherin 1, type 1 | 90.84 | $3.62 \times 10^{-14}$ | $5.54 \times 10^{-11}$ |
| NM_001300887 | AP1M2 | adaptor-related protein complex 1, mu 2 subunit | 68.77 | $1.71 \times 10^{-14}$ | $4.59 \times 10^{-11}$ |
| NM_020672 | S100A14 | S100 calcium binding protein A14 | 67.63 | $8.18 \times 10^{-13}$ | $6.50 \times 10^{-10}$ |
| NM_001260489 | LSR | lipolysis stimulated lipoprotein receptor | 65.26 | $1.09 \times 10^{-13}$ | $1.11 \times 10^{-10}$ |
| NM_016267 | VGLL1 | vestigial-like family member 1 | 61.44 | $5.20 \times 10^{-14}$ | $6.97 \times 10^{-11}$ |
| NM_001793 | CDH3 | cadherin 3, type 1, P-cadherin (placental) | 46.15 | $3.30 \times 10^{-12}$ | $1.97 \times 10^{-09}$ |
| NM_001010872 | FAM83B | family with sequence similarity 83, member B | 44.22 | $3.04 \times 10^{-14}$ | $5.40 \times 10^{-11}$ |
| NM_001104548 | MIR205HG;<br>MIR205 | MIR205 host gene; microRNA 205 | 41.9 | $7.06 \times 10^{-14}$ | $8.41 \times 10^{-11}$ |
| NM_005213 | CSTA | cystatin A (stefin A) | 35.73 | $1.36 \times 10^{-11}$ | $6.19 \times 10^{-09}$ |
| NM_001114978 | TP63 | tumor protein p63 | 33.42 | $1.22 \times 10^{-11}$ | $5.67 \times 10^{-09}$ |
| NM_003222 | TFAP2C | transcription factor AP-2 gamma | 28.75 | $2.70 \times 10^{-12}$ | $1.70 \times 10^{-09}$ |
| NM_174911 | FAM84B | family with sequence similarity 84, member B | 26.08 | $2.64 \times 10^{-13}$ | $2.36 \times 10^{-10}$ |
| NM_001171195 | ELAVL2 | ELAV like neuron-specific RNA binding protein 2 | 24.27 | $2.84 \times 10^{-11}$ | $1.17 \times 10^{-08}$ |
| NM_005555 | KRT6B | keratin 6B, type II | 22.52 | $7.27 \times 10^{-12}$ | $3.63 \times 10^{-09}$ |
| NM_001003407 | ABLIM1 | actin binding LIM protein 1 | 22.37 | $1.33 \times 10^{-12}$ | $9.62 \times 10^{-10}$ |
| NM_004482 | GALNT3 | polypeptide N-acetylgalactosaminyltransferase 3 | 20.14 | $3.31 \times 10^{-13}$ | $2.84 \times 10^{-10}$ |
| NM_001284406 | TMEM40 | transmembrane protein 40 | 17.94 | $2.38 \times 10^{-11}$ | $1.04 \times 10^{-08}$ |
| NM_001025252 | TPD52 | tumor protein D52 | 15.2 | 6.55 x 10<sup>-12</sup> | 3.48 x 10<sup>-09</sup> |
| NM_030949 | PPP1R14C | protein phosphatase 1, regulatory (inhibitor) subunit 14C | 12.83 | 1.11 x 10<sup>-12</sup> | 8.49 x 10<sup>-10</sup> |
| NM_001017970 | TMEM30B | transmembrane protein 30B | 11.99 | 1.83 x 10<sup>-11</sup> | 8.17 x 10<sup>-09</sup> |
| NM_001100812 | CXCL16 | chemokine (C-X-C motif) ligand 16 | 11.1 | 2.89 x 10<sup>-12</sup> | 1.77 x 10<sup>-09</sup> |
| NM_001127213 | LY6E | lymphocyte antigen 6 complex, locus E | 9.62 | 3.27 x 10<sup>-14</sup> | 5.40 x 10<sup>-11</sup> |
| NM_003105 | SORL1 | sortilin-related receptor, L(DLR class) A repeats containing | 7.79 | 1.10 x 10<sup>-11</sup> | 5.25 x 10<sup>-09</sup> |

This global analysis revealed that the epithelial/mesenchymal status of the exposed cells determines the entry of metallic NPs, althoughthe precise pathway remains unclear. Thus in order to gain more insight into the common genes involved in NP uptake, we investigated the pathways and functions attributed to the 541 genes. Data mining was performed using a GSEA analysis. **Table 4** reports the Hallmarks categories obtained from our data. Estrogen response (early and late) and KRAS signaling are more expressed in epithelial cells, whereas an EMT signature and hypoxia features are found in mesenchymal cells. The analysis based on REACTOME shows an enrichment in keratinization and cornification features, both indicators of epithelial cells, while mesenchymal associated categories concern extracellular matrix organization, glycosylation, and glycosaminoglycans (**Table 5**).

**Table 4:** Hallmarks enriched in epithelial and mesenchymal cells. Hallmarks that are mentioned all present a FDR-corrected q-value <0.05.

| HALLMARKS | SIZE | ES | NES | FDR q-val |
| --- | --- | --- | --- | --- |
| *Upregulated in Epithelial cells* |  |  |  |  |
| ESTROGEN_RESPONSE_LATE | 32 | -0.53 | -2.11 | 0.003 |
| KRAS_SIGNALING_UP | 16 | -0.60 | -1.97 | 0.004 |
| ESTROGEN_RESPONSE_EARLY | 32 | -0.41 | -1.67 | 0.027 |
| *Upregulated in Mesenchymal cells* |  |  |  |  |
| EPITHELIAL_MESENCHYMAL_TRANSITION | 64 | 0.60 | 2.86 | 0.000 |
| HYPOXIA | 17 | 0.70 | 2.37 | 0.000 |
| UV_RESPONSE_DN | 16 | 0.69 | 2.35 | 0.000 |

**Table 5:** REACTOME categories enriched in epithelial and mesenchymal cells. Categories that are mentioned all present a FDR-corrected q-value <0.05.

| REACTOME CATEGORY | SIZE | ES | NES | FDR q-val |
| --- | --- | --- | --- | --- |
| *Upregulated in Epithelial cells* |  |  |  |  |
| KERATINIZATION | 21 | -0.63 | -2.30 | 0.001 |
| FORMATION OF THE CORNIFIED ENVELOPE | 21 | -0.63 | -2.29 | 0.000 |
| *Upregulated in Mesenchymal cells* |  |  |  |  |
| EXTRACELLULAR MATRIX ORGANIZATION | 47 | 0.50 | 2.26 | 0.003 |
| DISEASES OF GLYCOSYLATION | 12 | 0.68 | 2.08 | 0.012 |
| DISEASES OF METABOLISM | 13 | 0.65 | 2.07 | 0.009 |
| NERVOUS SYSTEM DEVELOPMENT | 32 | 0.48 | 1.93 | 0.033 |
| GLYCOSAMINOGLYCAN METABOLISM | 11 | 0.64 | 1.88 | 0.041 |
| A TETRASACCHARIDE LINKER SEQUENCE IS REQUIRED FOR GAG SYNTHESIS | 8 | 0.70 | 1.86 | 0.041 |
| HEPARAN SULFATE HEPARIN HS GAG METABOLISM | 8 | 0.70 | 1.85 | 0.036 |
| METABOLISM OF CARBOHYDRATES | 13 | 0.59 | 1.84 | 0.035 |
| NCAM SIGNALING FOR NEURITE OUT GROWTH | 8 | 0.68 | 1.81 | 0.044 |
| ELASTIC FIBRE FORMATION | 12 | 0.59 | 1.81 | 0.041 |
| ECM PROTEOGLYCANS | 18 | 0.52 | 1.78 | 0.042 |

The comparison of our data with Chemical and Genetic perturbations database reveals an enrichment in signatures from breast cancer and epithelial to mesenchymal transition (**Figure 3A and Table 6**). For example, the ONDER_CDH1_TARGETS signature (down in E cells, and up in M cells) comes from a study by Onder *et al*^22^, where CDH1 expression was inhibited in HMLER cells (HMLE cells transformed with Ras oncogene), inducing an epithelial-mesenchymal transition. Thus, our data fit well with this HMLER cellular model used for understanding mechanisms governing the EMT program. Similarly, Charafe-Jauffret *et al* have produced signatures corresponding to differential expression between luminal, basal and mesenchymal cells^23^. Our data reveal luminal/basal features in E cells and mesenchymal ones in M cells. The stem cell signature by Boquest *et al*^24^ and Lim *et al*^25^ follows the same pattern, with a higher expression of corresponding genes in M cells compared to the E cells. In addition to a mesenchymal phenotype, Bontemps *et al* recently described the stem cells properties of HMLE-M cells^26^ .

**Figure 3:**
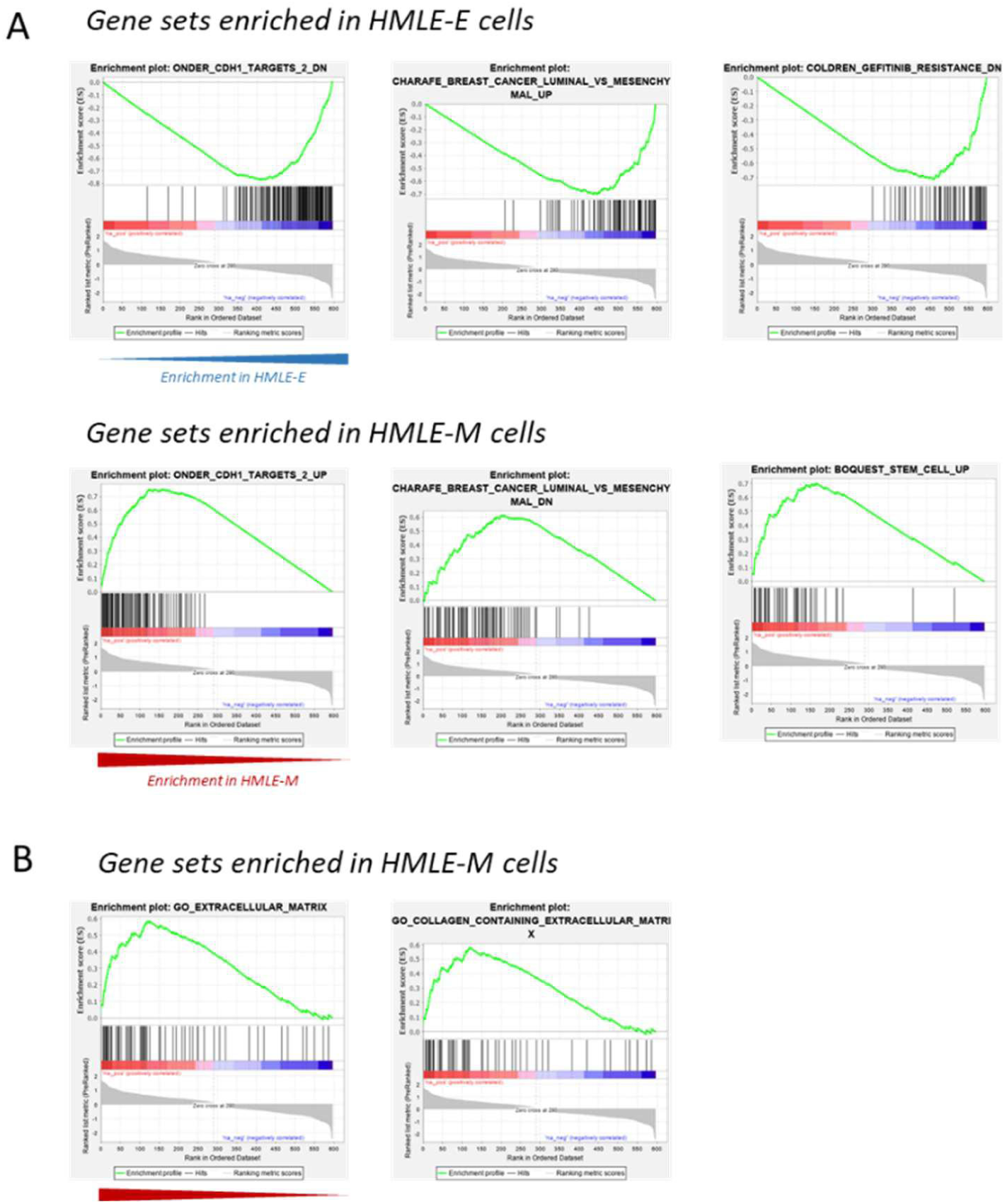
Gene Set Enrichment Analysis of nanoparticle uptake by HMLE-E and HMLE-M cells. **A.** Enrichment plots after analysis comparing the transcriptome dataset with the Chemical and Genetic Perturbations database, grouping signatures retrieved from published studies. The plots from the upper panel present signatures upregulated in HMLE-epithelial cells, whereas the lower panel presents genesets enriched in HMLE mesenchymal cells. **B.** Enrichment plots after analysis comparing the transcriptome dataset with the Gene Ontology database, Cell Component category. In HMLE mesenchymal cells, « Extracellular Matrix » and « Collagen-containing Extracellular Matrix » are the most enriched categories.

**Table 6:** Chemical and Genetic Perturbations categories enriched in epithelial and mesenchymal cells. Categories that are mentionned all present a FDR-corrected q-value <0.05.

| CHEMICAL AND GENETIC PERTURBATIONS | SIZE | ES | NES | FDR q-val |
| --- | --- | --- | --- | --- |
| *Upregulated in Epithelial cells* |  |  |  |  |
| ONDER_CDH1_TARGETS_2_DN | 145 | -0.77 | -4.30 | 0 |
| CHARAFE_BREAST_CANCER_LUMINAL_VS_MESENCHYMAL_UP | 73 | -0.70 | -3.50 | 0 |
| CHARAFE_BREAST_CANCER_BASAL_VS_MESENCHYMAL_UP | 45 | -0.78 | -3.39 | 0 |
| COLDREN_GEFITINIB_RESISTANCE_DN | 59 | -0.72 | -3.35 | 0 |
| HOLLERN_EMT_BREAST_TUMOR_DN | 45 | -0.73 | -3.23 | 0 |
| JAEGER_METASTASIS_DN | 64 | -0.64 | -3.10 | 0 |
| WAMUNYOKOLI_OVARIAN_CANCER_GRADES_1_2_UP | 22 | -0.75 | -2.77 | 0 |
| WAMUNYOKOLI_OVARIAN_CANCER_LMP_UP | 34 | -0.64 | -2.65 | 0 |
| LIM_MAMMARY_STEM_CELL_DN | 37 | -0.61 | -2.60 | 0 |
| AIGNER_ZEB1_TARGETS | 22 | -0.68 | -2.49 | 0 |
| *Upregulated in Mesenchymal cells* |  |  |  |  |
| ONDER_CDH1_TARGETS_2_UP | 91 | 0.75 | 3.94 | 0 |
| BOQUEST_STEM_CELL_UP | 45 | 0.70 | 3.11 | 0 |
| CHARAFE_BREAST_CANCER_LUMINAL_VS_MESENCHYMAL_DN | 78 | 0.61 | 3.09 | 0 |
| SCHUETZ_BREAST_CANCER_DUCTAL_INVASIVE_UP | 65 | 0.62 | 3.04 | 0 |
| CHICAS_RB1_TARGETS_CONFLUENT | 81 | 0.55 | 2.83 | 0 |
| MIYAGAWA_TARGETS_OF_EWSR1_ETS_FUSIONS_DN | 38 | 0.64 | 2.76 | 0 |
| LINDGREN_BLADDER_CANCER_CLUSTER_2B | 52 | 0.60 | 2.74 | 0 |
| GU_PDEF_TARGETS_UP | 23 | 0.73 | 2.72 | 0 |
| PICCALUGA_ANGIOIMMUNOBLASTIC_LYMPHOMA_UP | 33 | 0.67 | 2.71 | 0 |
| CHICAS_RB1_TARGETS_GROWING | 27 | 0.67 | 2.60 | 0 |

Finally, Enrichment analysis was conducted on Gene Ontology categories (**Table 7**). For the Cell Component ontology analysis, our data indicate again a pattern of epithelial cells in negatively regulated genes (basolateral/apical parts, cell-cell junctions, etc…) whereas most signatures present in M cells are focused on Extracellular matrix (ECM) and synaptic space (**Figure 3B**). ECM is also found in Molecular Function class, where the top classes identified concern ECM structural components, glycosaminoglycan binding and ion binding (e.g. zinc and calcium). Finally, Biological Process confirms the epithelial pattern in E cells (epidermal development and differentiation, cell-cell junctions). M cells exhibited a TGFβ response signature and features related to organ development.

**Table 7:**
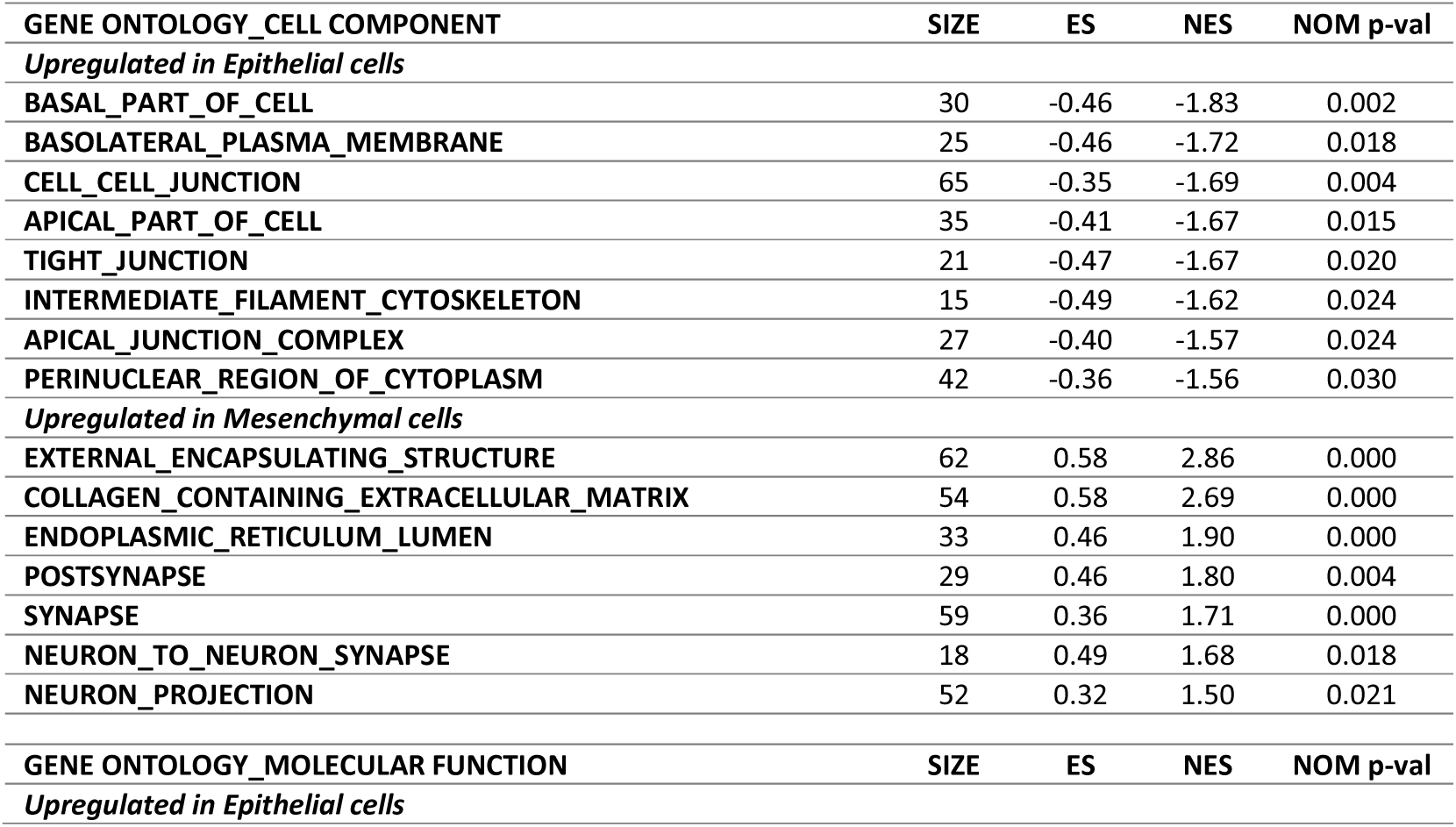
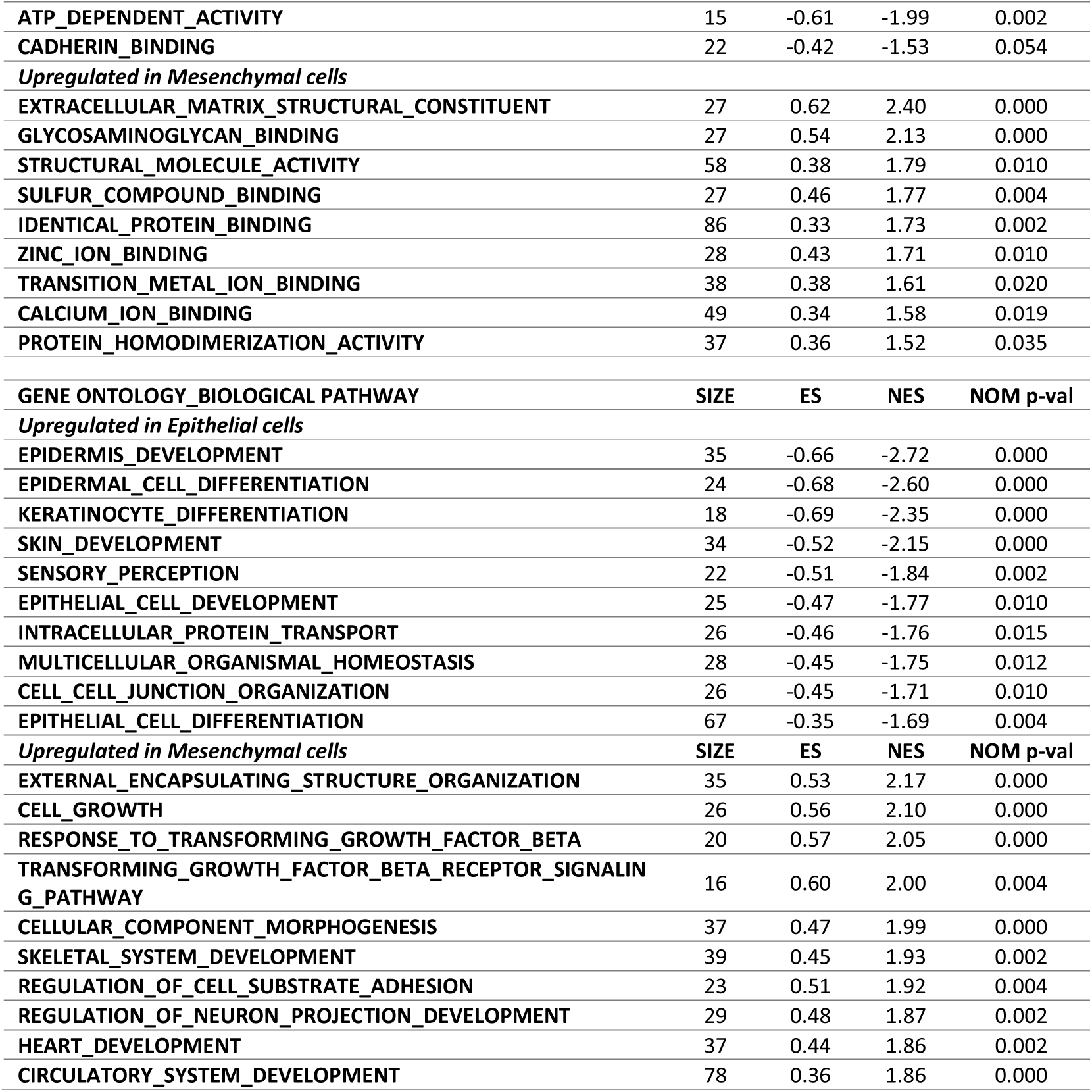
Gene Ontology categories enriched in epithelial and mesenchymal cells. Categories that are mentionned all present a nominal p-value <0.05 (except for cadherin binding)

Overall, these data suggest that typical epithelial and mesenchymal cell signatures are conserved upon nanoparticle exposure. However, among those features, there was a particular enrichment for ECM and cell membrane. Although this aspect seems obvious regarding the process of NP uptake, it indicates that the difference in uptake between E and M cells is mainly due to membrane and matrix differences between the cell types, and not to intracellular differences that could interfere. Indeed, one might have expected a large differential expression of intracellular vesicular transport or even transcription factors involved in E/M difference. The **supplementary table 2** details the differential expression data of genes belonging to the KEGG Endocytosis pathway. Interestingly, *DAB2* and *CLTCL1* are upregulated in mesenchymal cells exposed to NPs, suggesting a potential role for clathrin-mediated endocytosis in NP uptake. However, no specific receptor or binding molecule was identified by the analysis of these expression data, thus rather suggesting that the mechanism of uptake we were looking for should correspond to a marked difference in membrane or ECM between E and M cells.

One of the major differences between E and M cells at the membrane level is the expression of the CD44s molecule, which is highly specific and considered one of the most relevant molecular markers for distinguishing epithelial from mesenchymal cells in breast cancer. CD44 (Cluster of differentiation 44) is a glycosylated transmembrane protein encoded by 19 exons. The standard form (CD44s) consists of the first 5 exons corresponding to the hyaluronan-binding domain, and the 4 last exons of the locus encoding the cytoplasmic domain. Other isoforms are encoded with a combination of variable exons (v2 to v10). In the literature, the expression of CD44s is predominant in mesenchymal breast cell lines (MDA-MB-231, MDA-MB-436, MCF10A), while the V2-10 isoform is mostly expressed in more epithelial cells (MCF7, BT474, ZR-75)^27^.

*CD44* is not differentially expressed in the transcriptome data since the probes present on the microarray do not discriminate the *CD44s* isoform (specific for M cells) from the other *CD44* isoforms. However, CD44s is commonly used in breast cancer research, to identify and sort mesenchymal cells from patient tumours or *in vitro* models (e.g. with HMLE and MCF10A cells, **Figure 2A**). Hyaluronic acid (HA), one of the CD44 ligands, is synthesised by HAS (Hyaluronic Acid Synthases) enzymes^28^. In our data, *HAS2* is more expressed in M than in E cells (**Table 2**, FC=16.5, p-val=6.63×10^−12^), which could indicate a more important production of hyaluronic acid in mesenchymal cells, as was already described for breast cancer cells^29^. Taken together, these results indicate a possible upregulation of the CD44s/HAS2/HA axis in M cells.

### Evidence of the HA-CD44 axis role

To confirm the potential role of CD44-HAS2 in the uptake of NPs in our models, *CD44s* expression was measured in the 4 breast cancer cell lines, *i.e.,* T47D, MDA-MB-231, HMLE-E and HMLE-M. As expected, *CD44s* expression, the « standard » form of CD44, was found to be 10 times higher in HMLE-M than in HMLE-E, and 800 times more abundant in MDA-MB-231 cells compared to T47D (**Figure 4A**). Conversely, *CD44v2-10* isoform expression was 50-fold higher in the epithelial form of HMLE cells, and 6-fold higher in T47D compared to MDA-MB-231 cells (**Figure 4B**). In order to evaluate the potential direct role of CD44 in NP uptake, HMLE-M and MDA-MB-231 cells were transduced with lentiviral particles encoding a shRNA targeting *CD44*, with no isoform specificity. Epithelial cell lines were not included since the *CD44s* basal expression of these cell lines is very low. Reduced expression of *CD44s* and *CD44v* isoforms with the shRNA against CD44 was confirmed in both cell lines compared to a scramble shRNA coding vector (**Figure 4C**). The MDA-MB-231 cell line with a downregulation of *CD44* was further exposed to NM100 and NM103 TiO_2_ nanoparticles for 24 hours. The uptake of TiO_2_ NPs by the cells was evaluated by flow cytometry (**Figure 4D**) in three independent experiments, and replicated 3 additional times (**Supplementary Figure 4A and B**). Gold NPs uptake was also investigated (**Figure 4D**) and replicated (**Supplementary figure 4C**) to ensure that the results were not only linked to TiO_2_ chemical composition or large hydrodynamic sizes. Additionally, the **Supplementary Figure 5** shows the effect of CD44 expression inhibition on MCF10A-M cells. In all but one dataset, the uptake of TiO_2_ and gold NP was significantly reduced in cells transduced with the shRNA against CD44 compared to the scramble shRNA used as a control.

**Figure 4:**
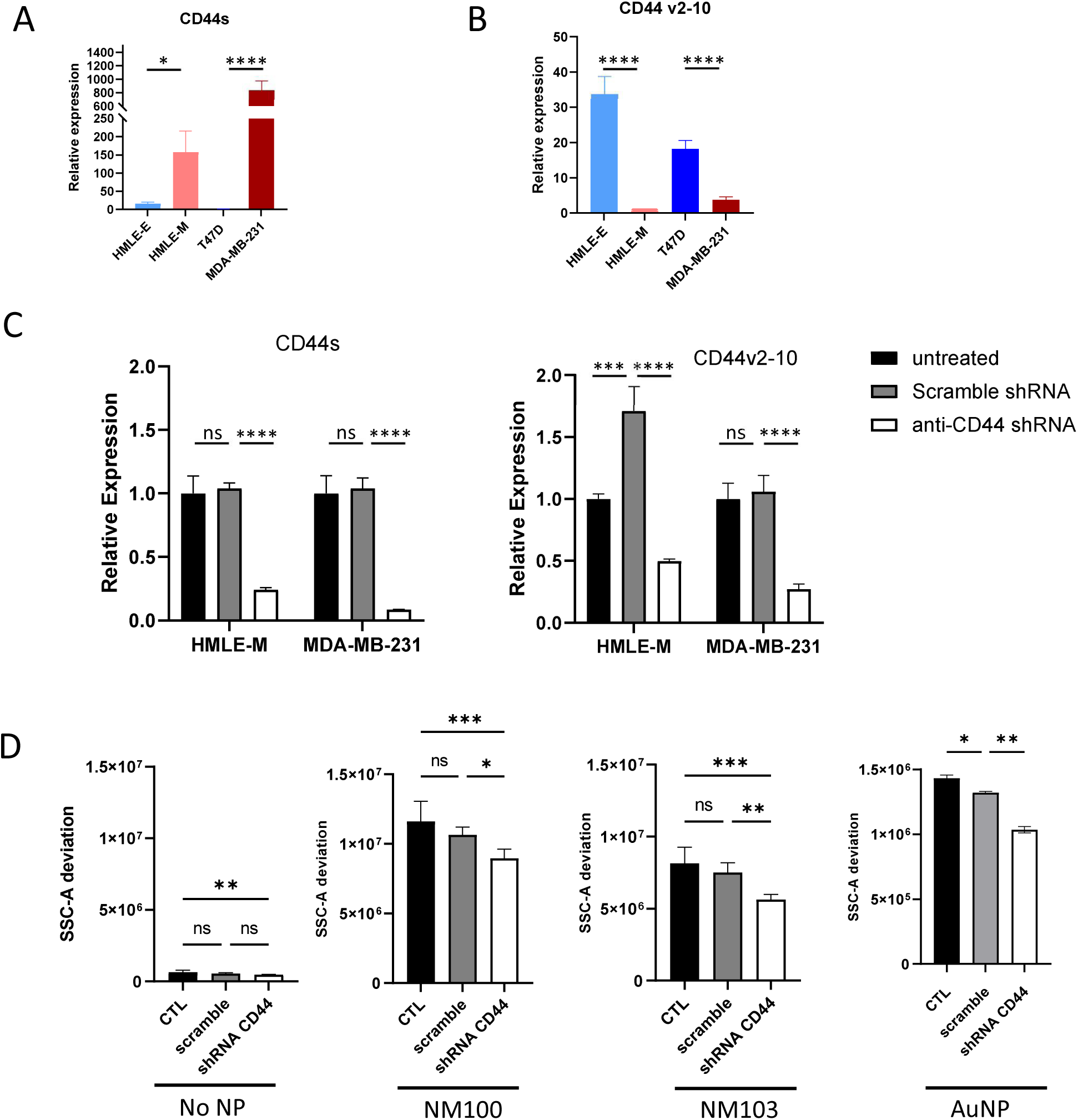
Expression levels of CD44 isoforms in different cell lines affect NP uptake. **A and B.** *CD44s* isoform (A) and *CD44v2-10* isoform (B) expression in 4 different breast cancer cell lines measured by qRT-PCR. Relative quantification was calculated with T47D (A) or HMLE-M (B) samples as reference samples. Measures of expression were performed in triplicates, and error bars represent RQ min and RQ max values. The statistical significance was determined by using the one-way Anova test, with a Sidak’s multiple comparison test. Note: ns, non-significant; * p < 0.05 and **** p < 0.0001. **C.** Quantification of *CD44s* and *CD44v2-10* expression in HMLE-M and MDA-MB231 cells after expression of shRNAs targeting all isoforms of CD44. Cells were transduced with lentiviral particles encoding anti-CD44 shRNAs and cultured for 9 days before RNA extraction. Relative quantification was calculated with untreated HMLE-M cells as a reference sample. Measures of expression were performed in triplicates, and error bars represent RQ min and RQ max values. The statistical significance was determined by using the one-way Anova test, with a Dunnett’s multiple comparison test. Note: ns, non-significant; *** p < 0.0005 and **** p < 0.0001. **D.** SSC-A deviation signals from MDA-MB231 cells expressing scramble or anti-CD44 shRNAs, unexposed to NPs (left panel), exposed to NM100 TiO_2_ NPs (middle panel), to NM103 TiO_2_ NPs (middle panel), or to AuNPs (right panel). The cells were exposed for 24 h to a NP concentration of 16 µg/cm² for TiO_2_ or 8 µg/cm² for AuNPs, and suspensions were subjected to flow cytometry analysis. The graph represents the mean of data from 10000 cells, in triplicates for TiO_2_ and duplicates for AuNPs. Error bars are SDs. Statistical significance is validated by Ordinary One-way ANOVA. Ns=non-significant, *indicates a p-value<0.05, **indicates a p-value<0.01 and ***indicates a p-value<0.001.

In order to investigate the role of CD44 in other tissues, its expression was inhibited in A549 and prostate models. Both A549 and A549-M cell lines could be modified, leading to a decrease in CD44 expression as confirmed by flow cytometry analysis in shRNA-CD44 lines (**Supplementary Figure 6A**). In the prostate model, only the WPE1-NB26 cell line could be modified as can be seen on **Supplementary Figure 7A**. We further exposed all cell lines to NM100, NM103 and gold NPs (**Supplementary Figure 6B and 7B**). While no effect of CD44 modification was evidenced in prostate, a significant decrease of AuNP endocytosis was shown in both lung lines. Therefore, CD44 could also be mediating metallic NP uptake in lung cells.

Hyaluronic Acid Synthases (*HAS1*, *2* and *3*) control the production and release of HA of different sizes on the extracellular side of the membrane^28^. HA is a high molecular weight molecule that binds notably to the extracellular domain of CD44. As *HAS1* expression was very low in all cell lines tested, we did not investigate the expression difference further. *HAS2* shows a clear differential pattern, with low to absent expression in epithelial lines (T47D, HMLE-E) and a much higher expression (>100-fold difference) in mesenchymal cell lines (**Figure 5A**). On the other hand, *HAS3* was expressed in all cell lines, with a higher expression in HMLE-E and T47D than in HMLE-M and MDA-MB-231 cell lines (**Figure 5B**). However, HAS3 was not differentially expressed in the transcriptome data of NP-exposed cells, thus reinforcing the choice of HAS2 as a good candidate for further investigation.

**Figure 5:**
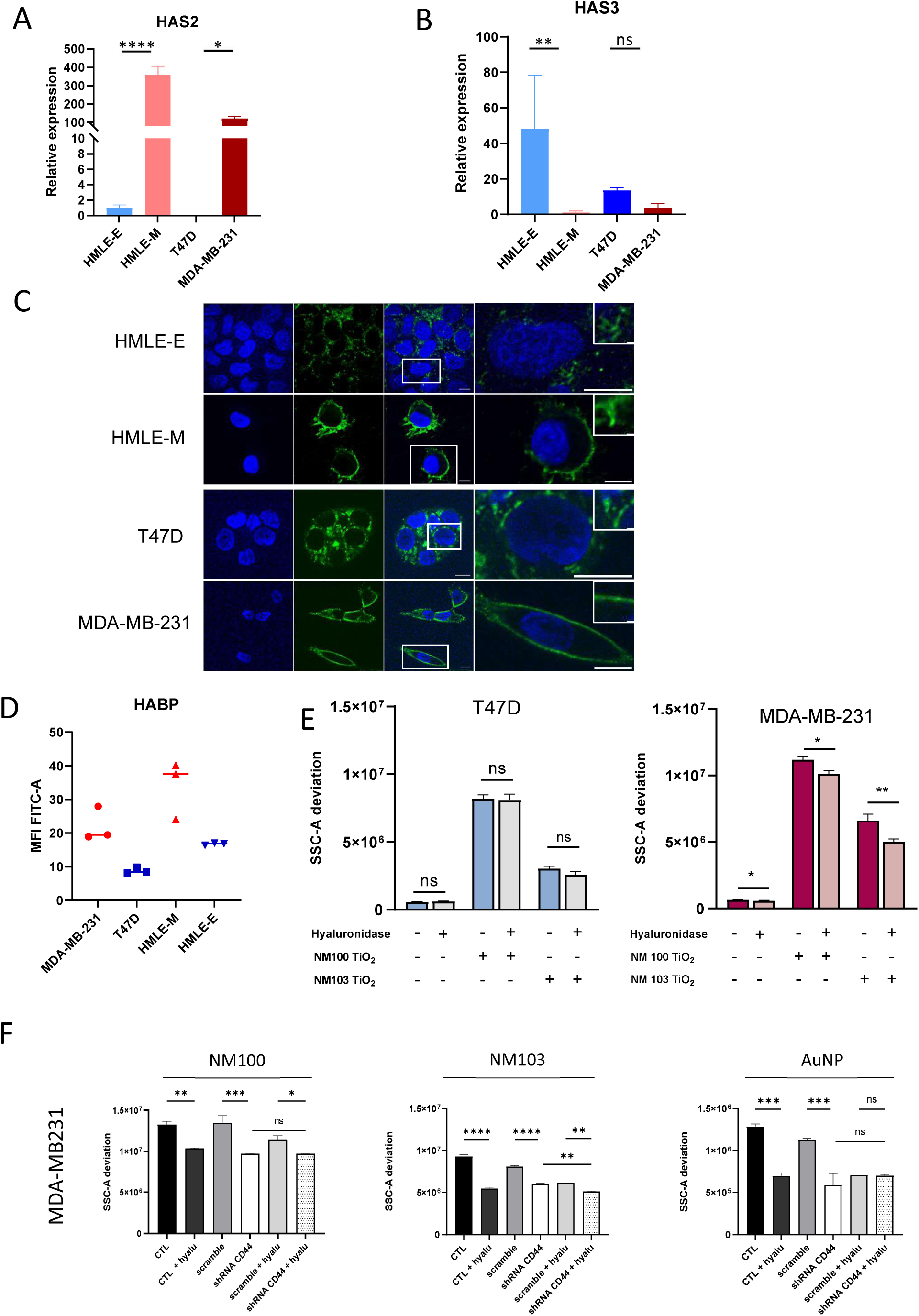
HAS2 expression, and effect of HA coat modulation on NP uptake. **A.** HAS2 expression in 4 different breast cancer cell lines measured by qRT-PCR. Relative quantification was calculated with HMLE-E sample as a reference. Measures of expression were performed in triplicates, and error bars represent RQ min and RQ max values. **B.** HAS3 expression in 4 different breast cancer cell lines, measured by qRT-PCR. Relative quantification was calculated with HMLE-M sample as a reference. Measures of expression were performed in triplicates, and error bars represent RQ min and RQ max values. For A and B, **t**he statistical significance was determined by using the one-way Anova test, with a Sidak’s multiple comparison test. Note: ns, non-significant; * p < 0.05, ** p <0.005 and **** p < 0.0001. **C.** Representative images of hyaluronic acid staining in the 4 different cell lines. Nuclei were stained with Hoescht (blue) and HA Binding proteins coupled to biotin were further detected using streptavidin-FITC (green). Images were obtained by confocal microscopy. Scale bar is 10 µm. **D.** Mean Fluorescence Intensity of HABP from at least 10000 cells for each of the 4 indicated cell lines, analysed by flow cytometry. Analysis was performed in triplicates. **E.** SSC-A deviation signals from T47D and MDA-MB231 cells treated with hyaluronidase and exposed to NM100, NM103 TiO_2_ or gold NPs. The graph represents the mean of data from 10000 cells, and in duplicates, error bars are SDs. The statistical analysis comparing untreated vs treated cells was performed with one-way anova. Ns=non-significant, *indicates a p-value<0.05 and **indicates a p-value<0.01. **F.** SSC-A deviation signals from MDA-MB231 cells expressing scramble or anti-CD44 shRNAs, treated with hyaluronidase and exposed to NM100 TiO_2_ NPs (left panel), to NM103 TiO_2_ NPs (middle panel) or AuNPs (right panel). The graph represents the mean of data from 10000 cells, and in duplicates, error bars are SDs. The statistical analysis comparing untreated vs treated cells was performed with one-way ANOVA and Sidak’s test. Ns=non-significant, *indicates a p-value<0,05 **p-value<0,01, ***p-value<0.001, ****p-value<0.0001

To assess the functionalityof HAS enzymes in the cell lines studied, we used fluorescence microscopy to detect Hyaluronic Acid Binding Protein (HABP) and thus estimate the HA levels at the cell surface. The cells were fixed and incubated with biotinylated HA-binding proteins, and further with fluorescently labeled streptavidin (**Figure 5C**). The microscopy analysis unveiled a distinct pattern of HA distribution, with a stronger signal at cell membranes in MDA-MB-231 cells compared to T47D cells. Similar results were obtained with the HMLE model. In agreement, quantification of HABP fluorescence signals by flow cytometry confirmed a more abundant HA labelling on the surface of mesenchymal cells compared to epithelial cells. (**Figure 5D**).

To validate the involvement of HA in NP uptake, HA content was modified by treating cells with hyaluronidase prior to exposure to nanoparticles (**Figure 5E**). In epithelial T47D cells, no effect of the hyaluronidase treatment was observed. In contrast, enzymatic digestion of HA on MDA-MB-231 cells significantly reduced NM100 and NM103 TiO_2_ NP uptake. A similar experiment was conducted on MDA-MB231 expressing the anti-CD44 shRNA (**Figure 5F**). In all three experiments (exposure to NM100, NM103 or AuNPs), the addition of hyaluronidase decreased significantly the uptake, confirming the results n Figure 5.E. The inhibition of CD44 expression also had a significant impact on internalization of the 3 NPs. The additive effect of inhibiting CD44 and digesting hyaluronan was seen only for NM103. Finally, TD7D, MDA-MB231 and HMLE cells were treated with 4-MU (4-methylumbelliferone), a hyaluronan synthesis inhibitor, before NP exposure **(Supplementary Figure 8**). A significant decrease in NM103 TiO_2_NP uptake can be observed in epithelial cells, but not in mesenchymal cell lines. However, caution should be exercised as 4-MU has been described as influencing breast cancer cells proliferation and phenotype^30^. Overall, these findings further confirm that HA coating facilitates NP uptake in mesenchymal cells, but may not be the only internalization route. All these data converge towards a model of opportunistic entry of metallic NPs into cells via their interaction with HA and endocytosis by CD44s.

### Metallic NP uptake via HA and CD44 is a general mechanism shared by mesenchymal breast cancer cell lines

In an attempt to generalize our observations to a larger panel of breast cancer cell lines, we measured *CD44s* RNA expression in 6 epithelial lines (T47D, SKBR3, BT20, MCF7, HMLE-E and MCF10A-E) and 5 mesenchymal lines (MDA-MB-231, MDA-MB-436, SUM159, HMLE-M and MCF10A-M). Consistently, *CD44s* expression was at least 10-fold higher in mesenchymal compared to epithelial cells (**Figure 6A**). Given that breast epithelial and mesenchymal cells are generally phenotyped by the CD24 and CD44 labelling, we performed flow cytometric phenotyping on 6 breast lines (not previously phenotyped, **Figure 6B**), confirming the elevated expression of CD44s in mesenchymal cells.

**Figure 6:**
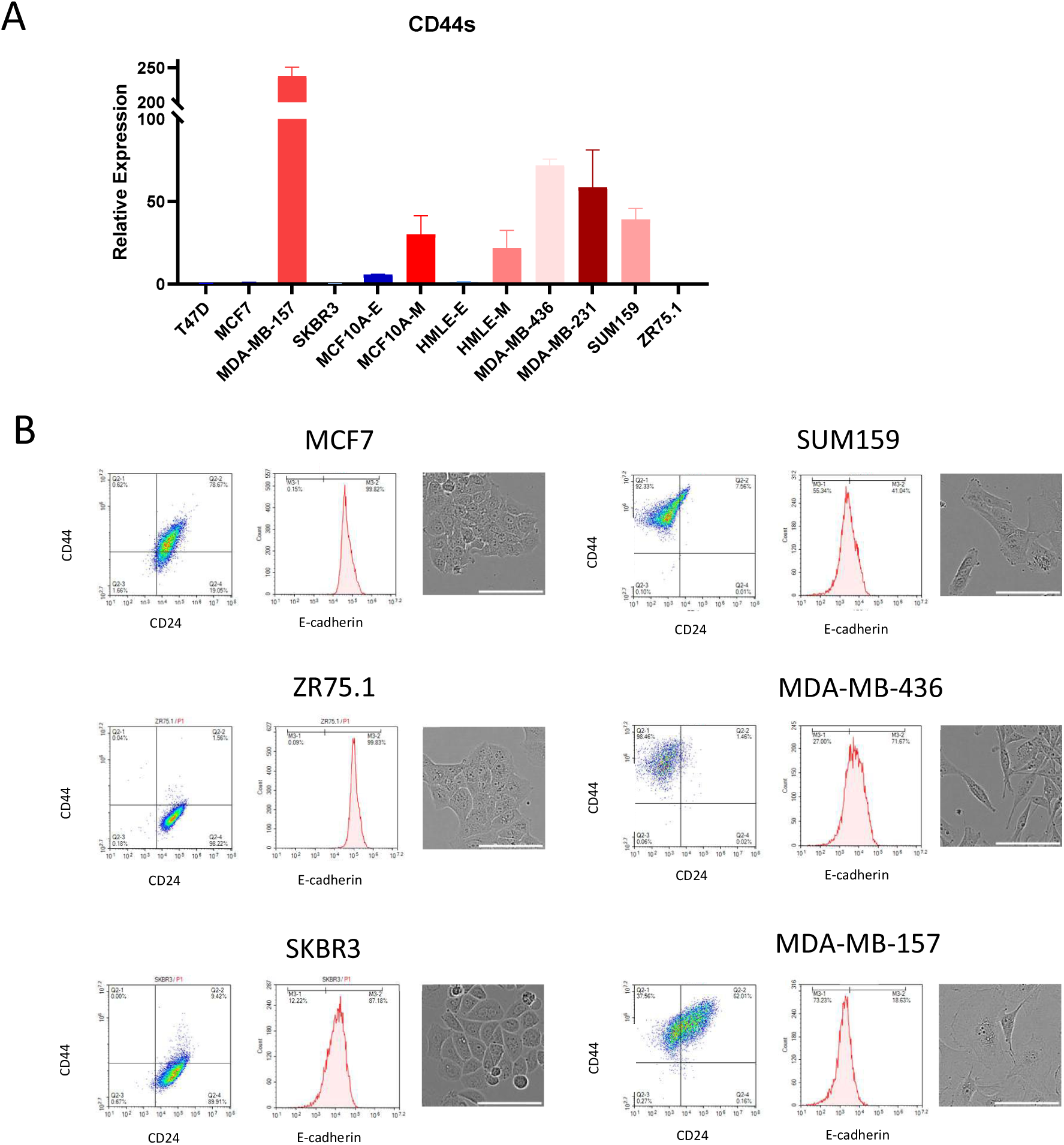
CD44s RNA and protein expression in a panel of breast cancer cell lines. **A.** CD44s expression measured by qRT-PCR in breast epithelial (T47D, SKBR3, ZR75-1, MCF7, HMLE-E, MCF10A-E, blue) and mesenchymal cell lines (MDA-MB-231, MDA-MB-436, MDA-MB-157, SUM159PT, HMLE-M, MCF10A-M, red). Measures of expression were performed in triplicates, and error bars represent RQ min and RQ max values. **B.** CD24/CD44 and e-cadherin phenotyping was performed by flow cytometry in epithelial and mesenchymal cells lines. The graph represents the mean of data from 10000 cells. The right panels show contrast-phase images of the 6 cell lines, scale bar 100 µm.

Next, *HAS2* expression was measured in the same cell lines, also indicating a higher quantity of *HAS2* mRNAs in mesenchymal models (**Figure 7A**). A labeling of HA with the HABP-streptavidin-biotin-FITC system confirmed by flow cytometry that HA was more abundant in mesenchymal cell lines compared to epithelial ones (**Figure 7B**). Next, fluorescence microscopy showed a stronger HABP signal at cell membrane in mesenchymal cells compared to epithelial lines (**Figure 7C**).

**Figure 7:**
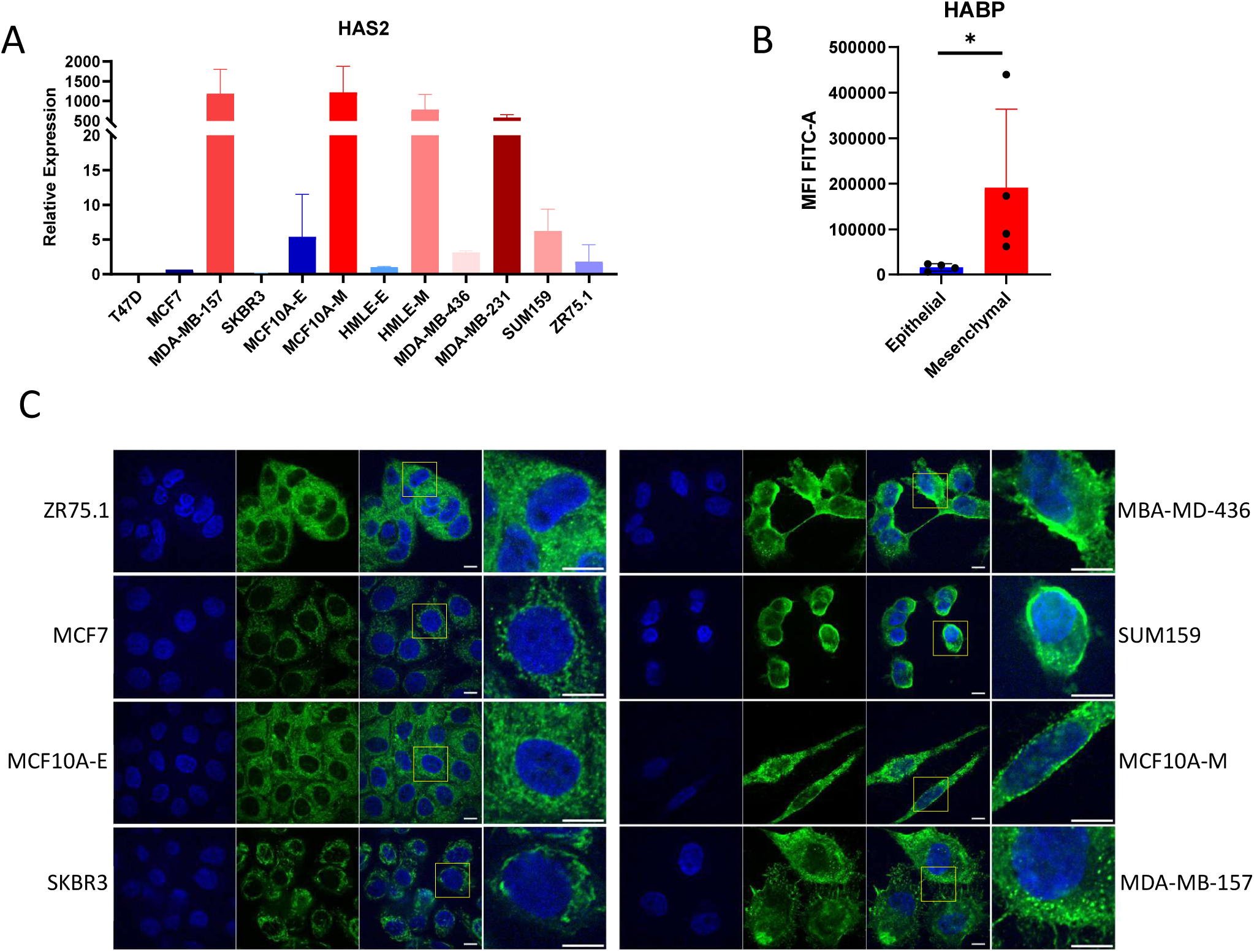
HAS2 RNA expression and HABP immunodetection in a panel of breast cancer cell lines. **A.** HAS2 expression measured by qRT-PCR in breast epithelial (T47D, SKBR3, ZR78-1, MCF7, HMLE-E, MCF10A-E) and mesenchymal cell lines (M231, M436, M157, SUM159PT, HMLE-M, MCF10A-M). Measures of expression were performed in triplicates and error bars represent RQ min and RQ max values. **B.** Mean Fluorescence Intensity of HABP-FITC in a mean of 10000 cells from each of the 8 cell lines, analysed by flow cytometry. Analysis was performed in triplicates. **C.** Representative images of hyaluronic acid staining in the 8 different cell lines. Nuclei were stained with Hoescht (blue) and HA Binding proteins coupled to biotin were detected with streptavidin-FITC (green). Images were obtained by confocal microscopy. Scale is 10 µm.

To determine the amount of NPs internalized in the cells from the panel of lines, all cell lines were exposed to 110 or 25 nm size TiO_2_ NPs, (NM100, **Figure 8A** and NM103, **Figure 8B**), and the SSC-A deviation was analysed after 24h of exposure. Despite some overlap, epithelial cell lines globally internalize less NPs than mesenchymal ones. Flow cytometry, as already mentioned, cannot be considered as a perfect method of quantification. Thus, absolute quantification of metal amount was measured with the ICP-MS method. Briefly, titanium and gold masses were quantified in cells exposed for 24h to TiO_2_ NPs (**Figure 8C**) or AuNPs (**Figure 8D**). For each type of NPs, the amount internalized in mesenchymal cells was higher than in epithelial cells, confirming again, the existence of a differential uptake linked to the phenotype of the cells.

**Figure 8:**
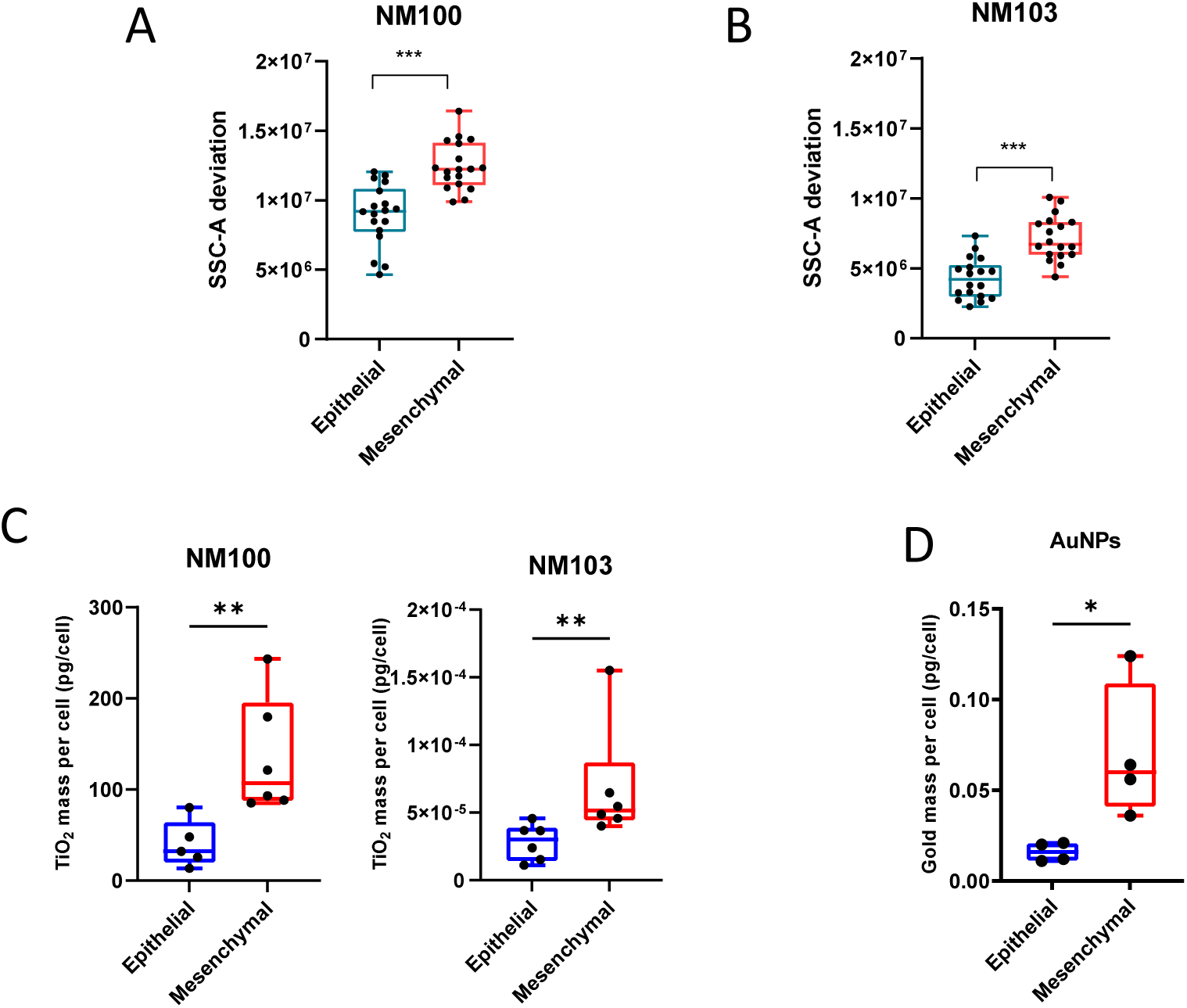
Nanoparticle uptake in breast cancer cell lines, according to the cell phenotype. **A.** SSC-A deviation of epithelial (blue) and mesenchymal (red) cell lines, after a 24 h exposure to NM100 TiO_2_ NPs at a concentration of 16 µg/cm². Each dot represents the mean deviation obtained with 10000 cells from each cell line. Statistical significance was evaluated with a Mann-Whitney test, *** indicates a p-value<0.001. **B.** SSC-A deviation of epithelial (blue) and mesenchymal (red) cell lines, after a 24 h exposure to NM103 TiO_2_ NPs at a concentration of 16 µg/cm². Each dot represents the mean deviation obtained with 10000 cells from each specific cell line. Statistical significance validated by a Mann-Whitney test, *** indicates a p-value<0.001. **C.** ICP-MS quantification of cytoplasmic titanium oxyde mass after a 24 h exposure with 16 µg/cm^2^ of NM100 or NM103 NPs, in 6 epithelial cell lines (T47D, MCF-7, HMLE.E, SKBR3, ZR75.1, MCF10A.E; blue) and 6 mesenchymal cells lines (MDA-MB-231, MDA-MB-157, HMLE.M, MDA-MB-436, SUM159PT, MCF10A.M; red). **D.** ICP-MS quantification of cytoplasmic gold mass after a 24 h exposure with 32 µg/cm^2^ AuNPs-PEG NH2 (32 nm), in 4 epithelial cell lines (T47D, MCF-7, HMLE.E, SKBR3; blue) and 4 mesenchymal cells lines (MDA-MB-231, MDA-MB-157, HMLE.M, MDA-MB-436; red). For C and D, **t**he statistical significance was determined by with a Mann-Whitney test. Note: * p < 0.05 and ** p <0.005.

## DISCUSSION

In addition to the physicochemical characteristics of NPs, cell parameters can have a dramatic impact on NP internalization. Cellular models and culture conditions such as cell density have been shown to influence uptake^31^. Other factors, such as the impact of the cell cycle phase, are more controversial and are at least partly due to changes in cell size as cells progress through the cell cycle. A simulation study has confirmed that the cell cycle does not play a role in NP uptake heterogeneity^32^. The genetic identity of cells also has an impact on endocytosis processes: for example, KRAS-mutant breast cancer cells have a higher NP uptake than wild-type cells, which is largely explained by a difference in macropinocytosis capacity of the cells^33^. In addition, the reliability and reproducibility of the results are also highly dependent on the methods used to study uptake. Indeed, cellular models, NP preparations, and analytical methods can be very heterogeneous from one study to another, and may introduce bias. One of the main methods used to characterize endocytic processes is the incubation of cells with specific inhibitors. However, several groups have cautioned about the lack of specificity of these inhibitors, their reduced activity in the presence of serum and the use of poorly standardized protocols often leading to misinterpretations^34,35^. We conducted toxicity assays that revealed the toxicity of many of the classical inhibitors (such as nocodazole, dynasore, and chlorpromazine) on our cellular models (data not shown), as already described by Sousa de Almeida *et al*^11^. Considering these limitations and in order to have a larger view of this complex question, we decided here to evaluate the uptake of different types of metallic nanoparticles in several independent cellular models and by using multiple complementary methodological approaches.

In this report as well as in previous research^17,18^, we noticed a systematic larger uptake of metallic NPs in mesenchymal compared to epithelial breast cancer cells. The different NP types covered a large range of hydrodynamic sizes (from 26 to 300 nm), negative and positive charges as well as various metallic components. To try to explain this difference in internalization and to avoid NP-linked biases, we performed a transcriptome analysis using four different NP suspensions. The analysis only included genes that were differentially expressed between epithelial and mesenchymal cells with a similar pattern among all NPs. As expected, most significant genes corresponded to epithelial-mesenchymal transition, but interestingly, the analysis revealed an enrichment in membrane-related phenomena and functions, suggesting that endocytosis could be the main process explaining the difference observed. The data did not allow the identification of a single major mechanism involving particular receptors as could be revealed in some studies^36^. Instead, it suggests a potentially more universal pathway used by non-decorated NPs, although not exclusive for NP internalization. Remarkably, the transcriptome analysis highlighted HAS2 as one of the most differential genes, and our data also showed a differential expression of CD44s between epithelial and mesenchymal cell lines (**Figures 4 and 6**). Finally, we could evidence the presence of hyaluronan at the cell surface of mesenchymal cells, while it appears mainly cytoplasmic in epithelial cells (**Figures 5 and 7**). Therefore, we focused on the hyaluronan-CD44 interaction, since hyaluronan synthesis and specific alternative splicing of CD44 are known markers of EMT in breast cancer.

While we were carrying out this study, Rodriguez team published two major studies showing that iron, copper and other metals could interact with hyaluronic acid, and further be internalized in cells via the recycling of HA bound to CD44^37,38^. Muller *et al.* showed that iron is internalized to a greater extent in mesenchymal HMLER CD44^high^ cells compared to more epithelial CD44 knock-out cells^38^. These observations are consistent with our hypothesis according to which metallic NPs would be taken up preferentially by cells with a more active HA-CD44 axis. A more recent study from the same group confirmed the involvement of this axis for other metals such as copper ^37^. In 2024, Tsai *et al* showed a preferential uptake of FePt NPs in mesenchymal compared to epithelial cells, through HA-CD44 as well^39^. They identified the link between CD44 expression, tyrosine kinase inhibitor resistance and mesenchymal state of cells and the endocytosis of FePt alloy NPs. Interestingly, they showed that this NP uptake was mediated by the binding of HABP2 protein to NP corona, therefore facilitating interactions with CD44. In addition, several groups have described a differential amount of hyaluronan between epithelial and mesenchymal cells, and Udabage *et al*^40^ have demonstrated that the daily turnover of HA is faster in MDA-MB-231 cells than in T47D cells. In this report we could demonstrate that upon treatment with inhibitors of the HA-CD44 axis, the endocytosis of several types of NPs was dramatically reduced, in several cell models (Figures 4 and 5). Although alterations in the composition of the protein corona, which may modulate NP uptake, were not examined in the present study, the broad range of NP core materials, sizes, and surface charges investigated across multiple breast cancer cell lines provides robust evidence supporting a conserved mechanism of metallic NP internalization. Our findings, together with existing literature, indicate a general mechanism for uptake of metallic nanoparticles and a link with EMT in breast cancer cells.

CD44 is also a marker of progression status in other tumors, such as ovary, prostate, lung or colon carcinoma^41^. We searched for a role of CD44 in NP uptake in two models of EMT in lung and prostate. While we could not link differential CD44 expression to a progression through EMT, a decrease in CD44 expression or a modulation of hyaluronan content had a significant impact on NP uptake. Complementary studies on relevant models of other tumor types and showing differential CD44 expression would be compelling.

A few studies including genome-wide approaches have been reported in the last years, interrogating corona proteins or cellular receptors involved in multiple types of nanoparticles endocytosis. Thus, LDL Receptor, SLC46A3 and SCARB1 were identified as receptors able to facilitate NP uptake within cellular models, either by binding known corona proteins^42^ or by regulating lysosome transport^43^. Interestingly, Montizaan *et al.*^36^ identified, in addition to the above-mentioned roles of LDLR and SCARB1, that glycosaminoglycans, and more precisely heparan sulfate metabolism, are also involved and provide a route for the internalization of NPs. Our study highlighted the role of hyaluronan, another glycosaminoglycan (GAG), in metallic NP uptake. Furthermore, our transcriptome analysis revealed that several GO categories corresponding to GAG metabolism (GAG_METABOLISM, Heparan_Sulfate METABOLISM, Chondroitin_sulfate _dermatan_sulfate METABOLISM) are found significantly enriched in mesenchymal cells compared to epithelial cells when exposed to NPs (Table 5).

Interestingly, other cellular models exhibit differential NP uptake. In the study of Pannico *et al*, both uptake kinetics and the amount of AuNPs were higher in prostate cancer cells compared to a non-cancerous epithelial cell line^44^. Guglielmi *et al* and Costanzo *et al* showed a higher uptake of PLGA (Poly lactic-co-glycolic acid) and mesoporous silica NPs in myoblasts compared to myotubes^45,46^, which represent more differentiated muscle. The authors postulate that this uptake difference could be due to the processes occurring along the differentiation. Another interesting plasticity phenotype is monocyte to macrophage differentiation and further macrophage polarization. McParland *et al* used PEGylated AuNPs to show a higher internalization in macrophages compared to monocytes, and a better uptake efficiency when macrophages were M2-type^47^. Taken together, these studies confirm an effect of differentiation in the NP uptake. Of course those processes (differentiation, polarization, EMT…) are accompanied by ECM changes, notably hyaluronan coat of cells^48^. Solier *et al* showed for example than inflammatory M1 macrophages were more prone to metal ions uptake, via the HA-CD44 axis, than primary monocytes^37^. It would be of great interest to perform a global survey of GAG or hyaluronan uptake processes in all those cell types and correlate this with NP internalization.

The higher uptake of nanoparticles in mesenchymal cells is an encouraging feature for nanomedicine approaches. Indeed, in most tumours, mesenchymal cells are the most aggressive, prone to metastasis and resistant to radio and chemotherapy. This is confirmed by the study of Brill-Karniely *et al*^49^, who showed a correlation between cancer aggressiveness features, such as metastatic capacity, cell uptake of particles, including nanoparticles, and cell deformability. In an elegant study, they also demonstrated that an EMT signature can be attributed to cells with the highest uptake capacity. In contrast, cells from the same cancer line, and selected for not uptaking NPs had a signature specific to epithelial cells. They could also In addition, several teams have successfully demonstrated the ability to identify drug-resistant and aggressive cells by sorting their capacity of NP uptake^50,51^. In another study, Ma *et al* showed that drug-resistant repopulating cells internalize more microparticles than original cells^52^. A 2024 study obtained a similar pattern using a lung carcinoma model, with a more efficient internalization in drug-tolerant persister cells compared to the original cell line^39^. All those research results can be related to the models used in our study. Indeed, the HMLE model, a relevant model for EMT process, harbours a higher NP uptake in mesenchymal cells that are radio- and chemoresistant and have extreme mesenchymal characteristics^19,26^.

Nonetheless, breast tumors are very complex and heterogeneous tissues that harbor epithelial and mesenchymal cells, with proportions that differ according to the histological subtype of the tumor. In addition, interactions with the extracellular matrix as well as other cell types (fibroblasts, immune cells) add another level of complexity. All these factors contribute to the challenges of designing effective anti-cancer treatments and the occurrence of relapse. Immune cells, notably monocyte-macrophages, have a high propensity for uptaking nanoparticles. Thus, it would be interesting to evaluate, in a tumor context, how metallic nanoparticles distribute between epithelial and mesenchymal cancer cells, immune cells, or remain within the extracellular matrix. In addition, the recent reports describing the *in vivo* use of metallic nanoparticles in radiation enhancement procedures have shown a striking effect of anti-tumor immune response^5,53,54^. Thus, a study of NP uptake related to CD44 expression *in vivo* is needed.

## CONCLUSIONS

In summary, this work describes a general mechanism used by metallic nanoparticles to enter breast cancer cells, and correlates nanoparticle uptake with EMT status via CD44 expression. By using EMT models and cell lines of breast cancer, we demonstrated a preferential endocytosis of various NP (with a broad range of sizes, charges and composition) in mesenchymal cells compared to the epithelial cells. Among the characteristic features related to EMT and which were likely to modulate particle uptake, we identified the hyaluronan-CD44 axis as one of the main players of NP endocytosis. CD44 expression inhibition and hyaluronan enzymatic digestion both led to a marked decrease in NP internalization. This impact of CD44 on NP uptake could also be observed in a lung cell line. Our results provide the link between NP uptake, EMT status of the cells and CD44 expression, and are consistent with the literature where CD44 is a well-known marker of EMT status and has been shown to modulate NP uptake in breast and lung. An *in vivo* approach would be necessary to confirm this preferential distribution of metallic NPs towards aggressive and resistant cells in the tumor context but also to determine whether their endocytosis in immune cells could impact radio-therapeutic strategies.

## METHODS

### Cell culture

Sixteen cancer cell lines were used for this study, among which twelve were from breast, 2 from lung (A549 and A549-M) and 2 from prostate (RWPE1 and WPE1-NB26). All cells were routinely cultured at 37°C in a humidified atmosphere containing 5% CO_2_ and checked for the absence of mycoplasma (MycoAlert™ Mycoplasma Detection Kits, Lonza). MDA-MB-231 (ATCC number HTB-26), T47D (ATCC number HTB-133), MDA-MB-157 (ATCC number HTB-24), MDA-MB-436 (ATCC number HTB-130), ZR-75-1 (ATCC number CRL-1500), SKBR3 (ATCC number HTB-30), MCF7 (ATCC number HTB-22) were grown in Dulbecco’s Modified Eagle’s Medium (DMEM GlutaMAX, Thermo Scientific), supplemented with 10% (v/v) heat-inactivated fetal bovine serum (Thermo Scientific), 1 mM antibiotic-antimycotic (Invitrogen; Thermo Fisher Scientific). SUM159-PT was cultured in Dulbecco’s Modified Eagle Medium F12 (DMEM F12 GlutaMAX, Thermo Scientific) supplemented with 5% (v/v) heat-inactivated fetal bovine serum (Thermo Scientific), 1 mM antibiotic-antimycotic (Invitrogen; Thermo Fisher Scientific) and additionally supplemented with 10 µg/ml of insulin (I9278-5ML, Sigma-Aldrich) and 0,5 µg/ml of hydrocortisone (H4001, Sigma-Aldrich). The R.A. Weinberg lab (Whitehead Institute for Biomedical Research, MIT) generously provided the HMLE cell line^19^. Upon treatment with 10 ng/ml of TGFβ1 (580704, Biolegend), HMLE epithelial cells transitioned into mesenchymal cells within a few days. They were cultured in Dulbecco’s Modified Eagle Medium F12 (DMEM F12 GlutaMAX, Thermo Scientific) supplemented with 10% (v/v) heat-inactivated fetal bovine serum (Thermo Scientific), 1 mM antibiotic-antimycotic (Invitrogen; Thermo Fisher Scientific) and additionally supplemented with 10 µg/ml of insulin (I9278-5ML, Sigma-Aldrich), 10 ng/mL human epidermal growth factor (E9644, Sigma-Aldrich) and 0,5 µg/ml of hydrocortisone (H4001, Sigma-Aldrich). The HMLE-M cells were cultured in the same medium, completed with 5 ng/ml of TGFβ1 in the culture medium. Similarly, the MCF10A cell line (ATCC number CRL-10317) transitioned into mesenchymal cell upon treatment with 10 ng/ml of TGFβ1 within a few days. They were cultured in Dulbecco’s Modified Eagle Medium F12 (DMEM F12 GlutaMAX, Thermo Scientific) supplemented with 5% (v/v) horse serum (11510516, Gibco, Fisher Scientific), 1 mM antibiotic-antimycotic (Invitrogen; Thermo Fisher Scientific) and additionally supplemented with 10 µg/ml of insulin (I9278-5ML, Sigma-Aldrich), 20 ng/mL human epidermal growth factor (E9644, Sigma-Aldrich), 0,1 µg/ml of Cholera toxin (C8052, Sigma-Aldrich) and 0,5 µg/ml of hydrocortisone (H4001, Sigma-Aldrich). Again, the mesenchymal status was maintained with 10 ng/ml of TGFβ1 added to the culture medium. The A549 and A549-M lines (ATCC number CCL-185) were grown in Dulbecco’s Modified Eagle’s Medium (DMEM GlutaMAX, Thermo Scientific), supplemented with 10% (v/v) heat-inactivated fetal bovine serum (Thermo Scientific), 1 mM antibiotic-antimycotic (Invitrogen; Thermo Fisher Scientific). The A549 cell line transitioned into mesenchymal cells (A549-M) upon treatment with 2.5 ng/ml of TGFβ1 within a few days. The RWPE1 (ATCC number CRL-3607) and WPE1-NB26 (ATCC number CRL-2852) cell lines were grown in KSFM medium with L-glutamine, EGF, and bovine pituitary extract (17005-075, Thermo Fisher Scientific), supplemented with 1 mM antibiotic-antimycotic (Invitrogen; Thermo Fisher Scientific).

### Nanoparticles

Titanium dioxide nanoparticles were obtained from Joint Research Centre Nanomaterials repository (JRC). Suspensions of NM100 and NM103 were prepared according to the Nanogenotox protocol at the European scale. Approximately 15 mg of powder was weighed in a scintillation vial of 20 ml and pre-wet with 30 µl of Ethanol 96°. Suspensions were prepared at a concentration of 2,56 mg/ml in 0,05% BSA before dispersing by ultrasonic probe for 7’54 at 20 % amplitude in a 4°C bath. Suspensions were characterized by Dynamic Light Scattering (DLS) measurements using a Nanosizer ZS (Malvern) in back-scattering configuration (173°). AuNP@PEG-NH2 and AuNP@PEG-COOH Nanoparticles were synthetized according to Turkevich method at Institut de Chimie Physique (ICP) by Pr. Sicard’s team^55^, and previously characterized^18^. AuNP@PEG were obtained from Sigma (reference 741965) and characterized by DLS (Table 1). Before any exposure, nanoparticles suspensions are mixed thoroughly and added to the culture medium, not directly in the culture wells to avoid any artefact. Since nanoparticles are subject to sedimentation by gravity^56,57^, a change in culture media volume may drastically modify the number of NPs in contact with the cell layer. Therefore, as recommended in nanotoxicology studies^58,59^, concentrations are expressed as the mass of NPs per surface unit of the culture dish (µg/cm²) A surface concentration of 16 µg/cm² will be equivalent to 72 µg/ml in 6-well plates, 58 µg/ml in 12-well plates and 61 µg/ml in 24-well plates.

### Expression modification with lentiviral vectors

Lentiviral vectors were generated at the Genetic engineering and protein biochemistry platform of CEA/IRCM. 293T cells (ATCC CRL-3216) were plated in 175 cm² flasks with a density of 9000 cells per cm^2^ and cultured in DMEM (4,5 g/L glucose + GlutaMAX™ Supplement + pyruvate) + 1 % penicillin-streptomycin + 10 % FCS for 72 hours. Cells were co-transfected with packaging plasmids (70 µg of pCMV-VSV-G : Addgene #8454 and 100 µg of pCMVΔR8.91^60^) and 100 µg of expression vector using the calcium phosphate method. The generated DNA precipitate was added to fresh medium and incubated with previously plated cells. Upon 17 hours of incubation at 37°C with 5 % CO2, the medium was changed, and cells cultured for additional 48 hours. Then, the culture medium containing viral particles was filtered on 0,45 µm. The filtered suspension was centrifuged at 83000g for 90 min at 4°C. Finally, the pellet was suspended in 200 µl PBS and frozen at −80°C. Virus titration was performed by exposing 10^5^ 293T cells to viral particles in 24-well plates. Four serial 5-fold dilutions were tested for each lentivirus, starting with a volume of 0.25 µl of frozen initial suspension. After 72 hours of culture, cells were washed three times with PBS, harvested by flushing, and the percentage of fluorescent GFP+ cells was further determined by flow cytometry. The viral titer is based on the condition where less than 30% of cells are GFP+, according to the following formula: titer (particles per µl) = (number of plated cells x %age of GFP+ cells)/volume of virus in this condition. The lentiviral pTrip-MND-GFP-H1-shCTRL (HCV) was used as a control vector encoding a shRNA targeting HCV and the pTRIP-MND-GFP-H1-SanD1-sh3_CD44 construct was used to encode shRNA sequences targeting CD44 (generous gift from J Calvo, iRCM, CEA). Transduction lasted for 3 days at 37°C and 5% CO2 in presence of scramble and shRNA-CD44 vectors (MOI = 10). Transduced cells were washed three times using PBS and harvested. After one week of culture, cells were sorted on GFP signal on a FACSAria^TM^.

### Hyaluronidase treatment

After 24 h of culture, cells were treated with 200 U/mL of hyaluronidase (H1115000, Sigma-Aldrich) for 2 h, before the exposure to 16 µg/cm² of TiO_2_ nanoparticles (58 µg/ml of NPs in 12-well plates). Cells were collected 24 h after exposure to nanoparticles and washed twice in PBS. The SSC deviation signals were measured by flow cytometry, for 10000 cells in each sample. Statistical analysis was performed with GraphPad Prism 9.

### 4-methylumbelliferone

24h after plating, cells were treated with 1mM of 4-MU (M1381, Sigma-Aldrich) for 24 h, before the exposure to 16 µg/cm² of TiO_2_ nanoparticles or 8 µg/cm² of AuNPs. Cells were collected 24 h after exposure to nanoparticles and washed twice in PBS. The SSC deviation signals were measured by flow cytometry, for 10000 cells in each sample.

### Flow cytometry

Cells were harvested from flasks or plates using a 0.05% TrypLE express solution (Fisher Scientific, Illkirch, France).and a 5-minute centrifugation at 300 g. After two PBS washes, Cells were counted using an automated cell counter (TC20, Biorad, Marnes la Coquette, France), with trypan blue exclusion. A total of 200.000 cells were stained with 2 µl of each of the following antibodies: CD24-PE (ML5 clone, 562789, BD BioScience), CD44 FITC (555478 BD BioScience) or CD44-BV421 and e-cadhérine APC – CD324 (EXbio) for 15 min at 4°C. Cells were washed and centrifuged for 5 min at 300 g and pellets were resuspended in PBS-1% BSA. Cells were analysed on a Novocyte instrument (ACEA Biosciences), considering at least 10000 events per sample. Statistical analysis was performed with GraphPad Prism 9.

### Darkfield microscopy

Darkfield microscopy was used to visualize gold and TiO_2_ nanoparticles inside cells. Cells were plated cells on microscopic slides 24 h before exposure to nanoparticles. Ti0_2_ NPs (NM100 and NM103) were added at a concentration of 6,46 µg/cm² and AuNPs-NH2 were added at 14,3 µg/cm². After incubation, slides were washed twice and fixed using a 4% paraformaldehyde solution for 15 min. After rinsing with PBS, the slides were mounted with ProlonGold anti-fade reagent (11539306, Fisher Scientific.). The slides were analysed under a BX40 Olympus microscope, with a 50X objective.

### HAPB detection by confocal microscopy and flow cytometry

For detection with microscopy, cells were seeded on 4-well glass slides (80427, IBIDI). After 72 h of culture, cells were washed with PBS and fixed with a 4% paraformaldehyde solution for 15 min. A PBS washing was followed by an incubation with 2% BSA for 1 hour at room temperature. Cells were then incubated with biotinylated Hyaluronic Acid Binding Protein (HABP, ref 385911-50UG, Millipore) at a dilution of 1:250 in a blocking buffer for 1 h. The cells were rinsed 3 times with PBS before incubation with Streptavidin-Alexa Fluor^TM^ 488 goat anti-mouse secondary antibody (Invitrogen, ThermoFisher, Illkirch, France), at a dilution of 1:500 in a blocking buffer for 45 min. Cells were rinsed 3 times with washing buffer and stained with 1 μg/mL Hoechst 33342 for 15 min. After a final rinse, slides were mounted with ProlonGold anti-fade reagent (Fisher Scientific, Illkirch, France) and sealed. Stained cells were analysed on a NIKON Ti2 / GATACA, W1 spinning disk microscope with 60X oil immersion objective and images were treated using the Fiji cell-image analysis software.

For HAPB detection by flow cytometry, 200000 cells were harvested from culture supports, and were centrifuged for 5 min at 300 g. Pellets were resuspended in PBS-1% BSA with Biotinylated Hyaluronic acid binding protein (385911-50UG, Millipore) at a dilution of 1:125, for 90 min. After that, cells were washed twice and pellets were resuspended in PBS-1% BSA with streptavidin Alexa fluor 488 (JIR016-540-084, Ozyme) at a dilution of 1:1000 for 30 min. After another two washes, pellets were resuspended in PBS-1% BSA. Cells were analysed on a Novocyte instrument (Agilent). Statistical analysis was performed with GraphPad Prism 9.

### ICP-MS

Cells were seeded in 25 cm^2^ flasks and exposed to 32 µg/cm² of AuNP for 6 h and 24 h, or 16 µg/cm² of TiO_2_ NPs for 24h. 32 µg/cm² is equivalent to a concentration of 160 µg/ml in 25 cm² flasks, and 16 µg/cm² is equivalent to 72 µg/ml. Cells were washed three times in PBS, before being harvested. Cell counts were performed in triplicate using an automatic cell counter (TC20, Biorad, Marnes la Coquette, France), followed by centrifugation. For gold quantification, cell pellets were digested in a nitric acidsolution, according to the standard procedures of the LNE (Laboratoire National de Métrologie et d’Essais, Paris, France; www.lne.fr). For titanium quantification, cells were digested in hydrofluoric acid and nitric acid, according to LNE’s procedures. The total titanium or gold masses were measured using Inductively Coupled Plasma–Mass Spectrometer (ICP–MS) and normalized to the cell number in each sample. Three measures were performed per sample and analysed with GraphPad Prism 9.

### Transcriptome Analysis

For both HMLE-E and HMLE-M cells, 100000 cells were plated in 6 wells plates and cultured as described above. After 24 h, cells were exposed for 24 or 48 hours to 16 µg/cm² (equivalent to 72 µg/ml) TiO_2_ NM100, TiO_2_ NM103, AuNPs-COOH, or AuNPs-NH2 (table 1). Briefly, a 0.05% TrypLE express solution (Fisher Scientific, Illkirch, France) was added to each well and incubated at 37°C until dissociation, followed by a 5 minutes centrifugation at 300 g. Cell pellets were then snap-frozen and stored at −80°C.

Total RNA was extracted from frozen HMLE cell pellets with Total RNA purification kit (reference 17240, Norgen Biotek Corp.), according to the manufacturer’s instructions. DNAse treatment was performed using TURBO DNA-free kit (Invitrogen; Thermo Fisher Scientific, Inc.). After validation of the RNA quality with Bioanalyzer 2100 (using Agilent RNA6000 nano chip kit), 100 ng of total RNA were reverse-transcribed following the GeneChip® WT Plus Reagent Kit (Thermofisher). Briefly, the resulting double strand cDNA was used for *in vitro* transcription with T7 RNA polymerase (all these steps are included in the WT cDNA synthesis and amplification kit of Thermofischer). After purification according to Thermofisher protocol, 5.5 µg of Sens Target DNA were fragmented and biotin labelled. After control of fragmentation using Bioanalyzer 2100, cDNA was then hybridized to GeneChip® Clariom S Human (Thermofischer) at 45°C for 17 hours. After overnight hybridization, chips were washed on the fluidic station FS450 following specific protocols (Thermofischer) and scanned using the GCS3000 7G. The scanned images were analyzed with Expression Console software (Thermofischer) to obtain raw data (cel files) and metrics for Quality Controls. Raw data were normalized using the Robust Multichip Algorithm (RMA) in Bioconductor using Brain array EntrezGene CDF version 22 for annotations^61^. Statistics were performed using Partek GS 7.0 ®. Afterwards, gene expression and comparison between samples was analyzed thanks to the TAC software (Applied biosystems). ANOVA was used to detect genes with a gene-level fold change >2 or <-2, and with a FDR-corrected p-value <0.05.

### Expression analysis by RT-qPCR

Total RNA was extracted from cell pellets with the RNeasy Plus Micro kit (Qiagen) and reverse transcribed with random primers and Superscript IV VILO (Invitrogen), following the manufacturer’s recommendations. Quantitative PCR was performed using the iTaq™ Universal Probes Supermix (Bio-Rad) and Taqman probes (Thermofisher). The references of the Taqman probe sets used are as follows: GAPDH : Hs99999905_m1, *CD44v2-10* : Hs01075866_m1, *CD44s* : Hs01081473_m1, *HAS2* : Hs00193435_m1, *HAS3* : Hs00193436_m1. A ΔΔCt method was used for comparative analysis, using GAPDH as a control gene. A RQ calculation was performed as recommended, using appropriate control sample depending on the experiment. Statistical analysis was performed with GraphPad Prism 9.

## Supporting information

supporting info Hullo

## ASSOCIATED CONTENT

## Supporting Information

Supplementary figures are available. **Figure S1**: SSC-A deviation signals from HMLE-E and HMLE-M cells plated at different densities and exposed to NM103 TiO_2_. **Figure S2:** Uptake of TiO_2_ and gold nanoparticles in a lung model of Epithelial-to-mesenchymal transition (A549). **Figure S3:** Uptake of TiO_2_ and gold nanoparticles in two prostate cell lines. **Figure S4:** CD44 expression modification affects NP uptake in MDA-MB231 line (replicate experiments). **Figures S5, S6 and S7**: CD44 expression modification affects NP uptake in MCF10A-M, A549 and WPE1-NB26 lines. **Figure S8:** Effect of 4-methylumbelliferone on nanoparticles uptake in T47D and MDA-MB231 cells.

## AUTHOR INFORMATION

## Author Contributions

Performed experiments: MH, CM, NF, CL, GP, JN, EB; Analysed data : MH, EB; Wrote the manuscript : CM, MH, EB; Secured funding and supervised the study : AC, SC, EB. All authors have read and agreed to the published version of the manuscript.

## ACKNOWLEDGEMENTS

This work was supported by the Transverse Division n°4 (Radiobiology) of the French Alternative Energies and Atomic Energy Commission (CEA). M. Hullo was a recipient of a PhD fellowship from the French Ministry of Higher Education, Research and Innovation. The work was supported by the iNanoTheRad Interdisciplinary object of Paris-Saclay University. The authors thank Romain Grall (previously at CEA) for initial contribution to the project. The ICP–MS experiments were conducted on the Platform of the LNE laboratory (Laboratoire National de Métrologie et d’Essais, Paris, France). Creation and production of the lentivirus was performed by the Genetic engineering and protein biochemistry platform of CEA/IRCM (Fontenay-aux-Roses, France) and transduced cells were sorted by Nathalie Dechamps on the cytometry platform of CEA/IRCM (Fontenay-aux-Roses, France). The authors are very grateful to Julien Calvo from CEA who generously gave access to lentiviral products. We also thank Sébastien Jacques from the GENOMIC’ platform of Cochin Institute for the microarray experiments.

