## supporting info Hullo for "The uptake of metallic nanoparticles in breast cancer cell lines is modulated by the hyaluronan-CD44 axis"

**low**

**medium**

**high**

**Confluence level**

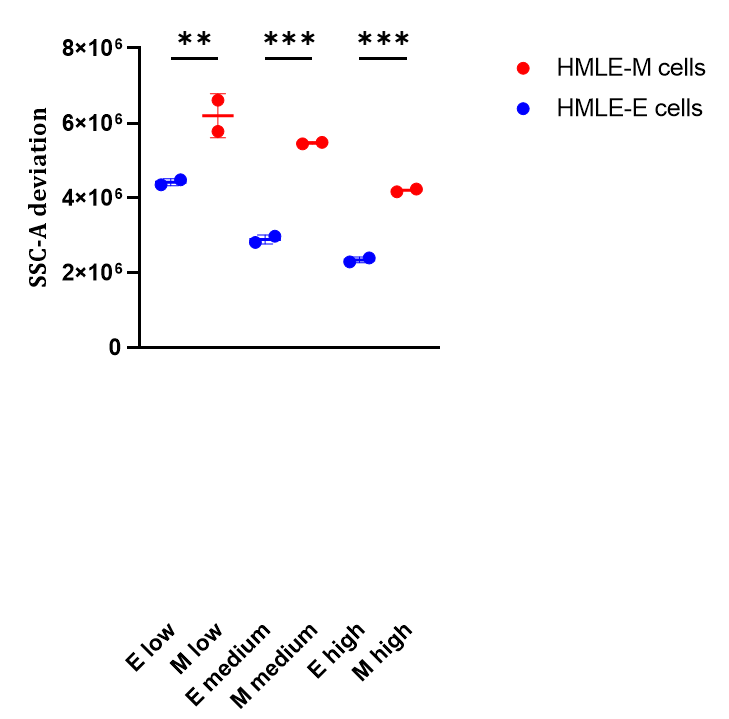

**Supplementary figure 1** : SSC-A deviation signals from HMLE-E and HMLE-M cells exposed to NM103 TiO_2_ NPs at a concentration of 10 µg/cm². Cells were plated in 24 wells plates at 3 different plating densities (10.000, 30.000 or 45.000 cells, corresponding respectively to low, medium and high levels of confluence) and exposed to NPs 24 h later. After 48 h exposure suspensions were subjected to flow cytometry analysis. The graph represents the mean of data from 10000 cells, analysed in duplicates. Errors bars are SDs. The statistical significance was determined by using an ordinary one-way Anova test, and Sidaks’ multiple comparisons test. Note: *** p < 0.0005 and ** p < 0.005.

**A549**

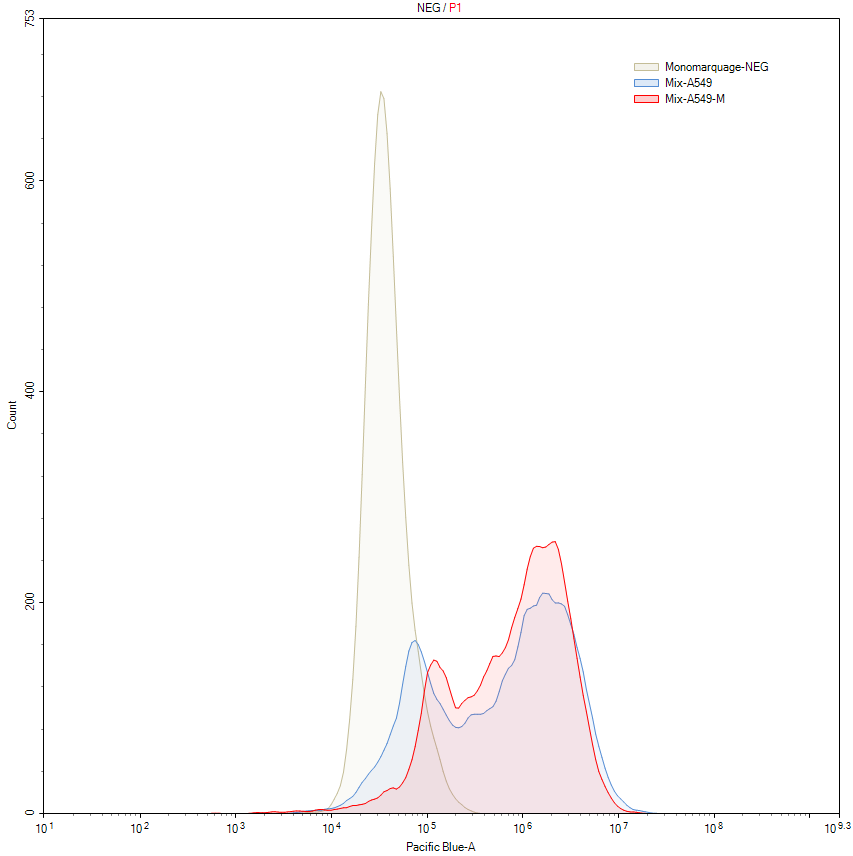

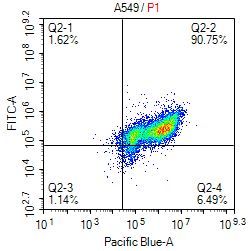

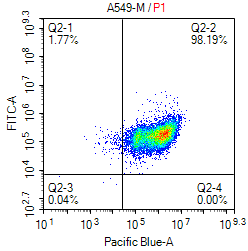

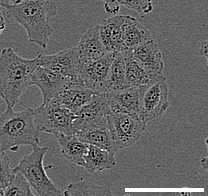

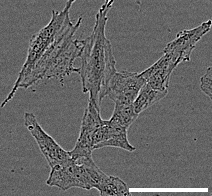

Count

CD44

Negative CTL

A549

A549-M

CD44

CD44

**A549-M**

+TGFβ

A

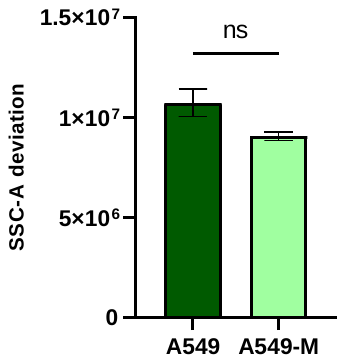

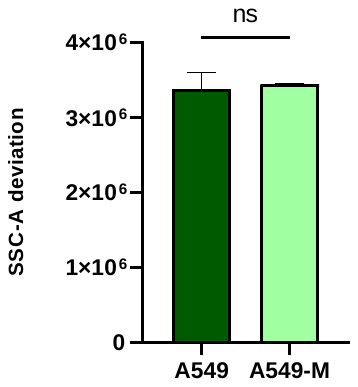

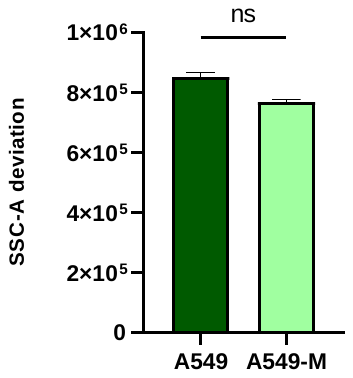

NM100

NM103

AuNP

B

**Supplementary Figure 2 :** Uptake of TiO_2_ and gold nanoparticles in a lung model of Epithelial-to-mesenchymal transition.

**A.** Model of induced epithelial to mesenchymal transition *in vitro* with TGFβ. A549 and A549-M cell lines were phenotyped by flow cytometry with CD44-BV421 (Pacific blue) antibodies (left panel). The middle panels show contrast-phase images of the 2 cell lines, magnification x10. Scale bar 100 µm. The right panel shows the overlay of CD44 expression between A549 and A549-M cells. **B.** SSC-A deviation signals from A549 and A549-M cells exposed to NM100 TiO_2_ NPs, NM103 TiO_2_ NPs and AuNPs at 16 µg/cm² for TiO_2_ and 8 µg/cm^2^ for AuNPs. Cells were exposed for 24 h, and suspensions were subjected to flow cytometry analysis. The graph represents the mean of data from 10000 cells, and in duplicates, error bars are SDs. The statistical significance was determined by using the Mann-Whitney test. Note: ns, non-significant.

**RWPE1**

NM100

NM103

AuNP

CD44

**WPE1-NB26**

A

B

Count

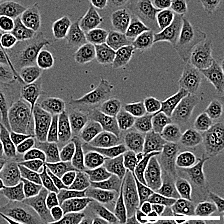

CD44

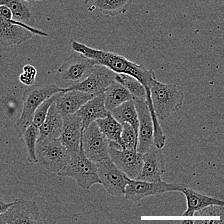

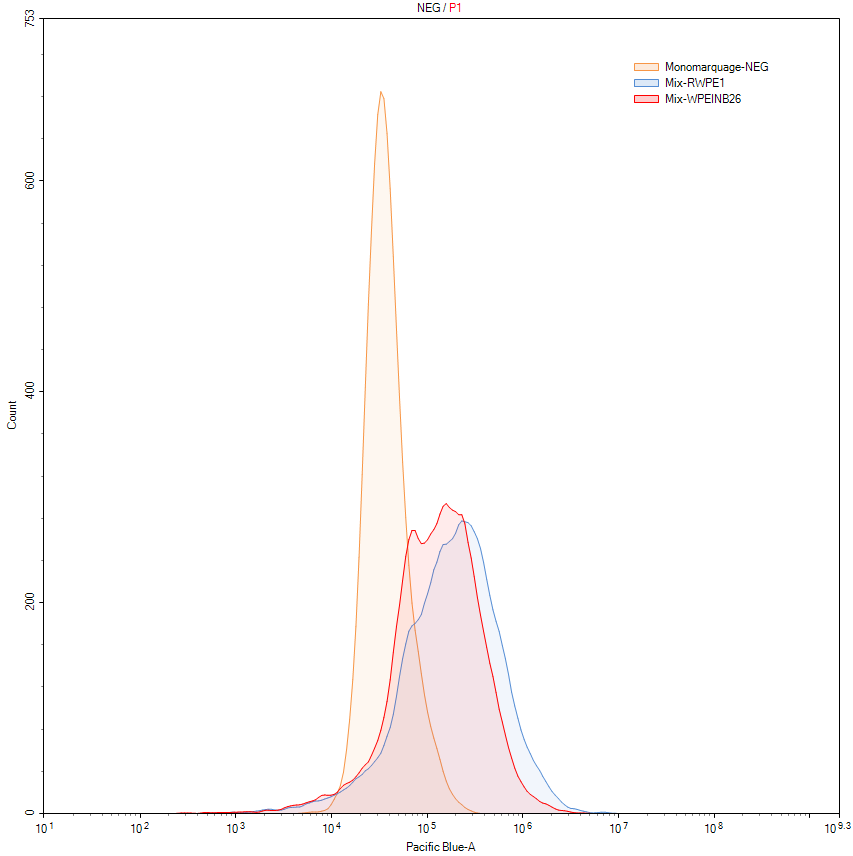

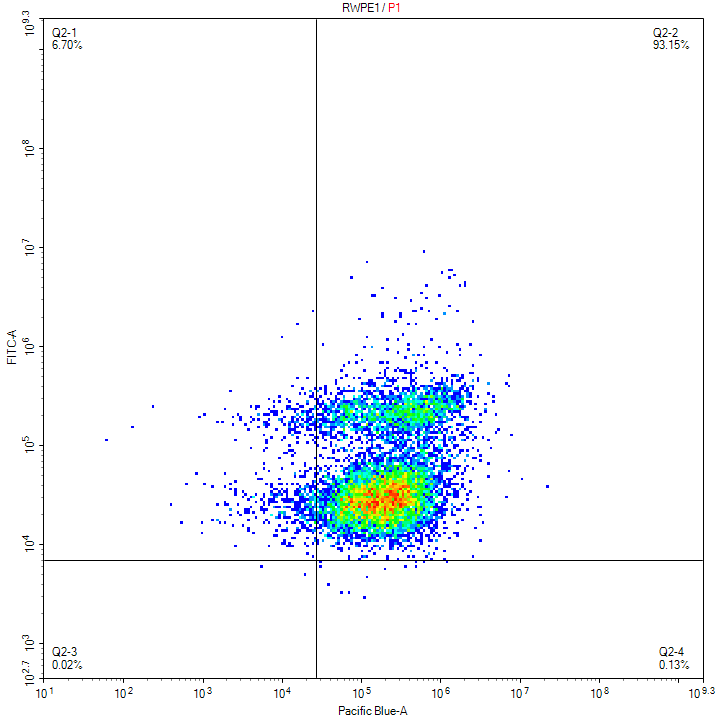

Negative CTL

RWPE1

WPE1-NB26

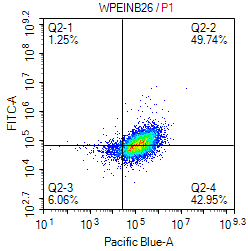

CD44

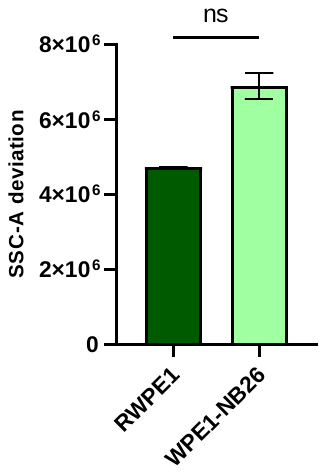

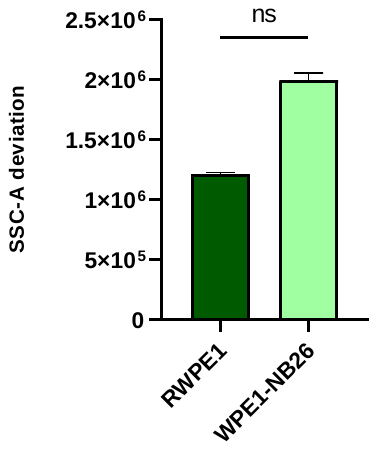

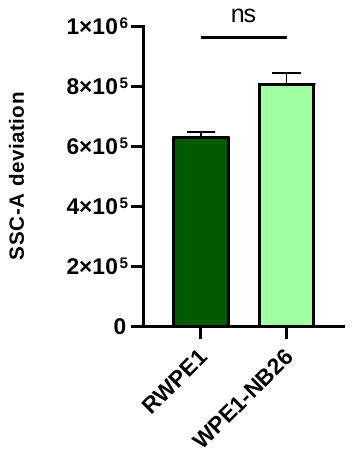

**Supplementary Figure 3 :** Uptake of TiO_2_ and gold nanoparticles in two prostate cell lines.

**A.** RWPE1 and WPE1-NB26 cell lines were phenotyped by flow cytometry with CD44-BV421 (Pacific blue) antibodies (left panel). The middle panels show contrast-phase images of the 2 cell lines, magnification x10. Scale bar 100 µm. The right panel shows the overlay of CD44 expression between RWPE1 and WPE1-NB26 cells. **B.** SSC-A deviation signals from RWPE1 and WPE1-NB26 cells exposed to NM100 TiO_2_ NPs, NM103 TiO_2_ NPs and AuNPs at 16 µg/cm² for TiO_2_ and 8 µg/cm^2^ for AuNPs. Cells were exposed for 24 h, and suspensions were subjected to flow cytometry analysis. The graph represents the mean of data from 10000 cells, and in duplicates, error bars are SDs. The statistical significance was determined by using the Mann-Whitney test. Note: ns, non-significant.

A

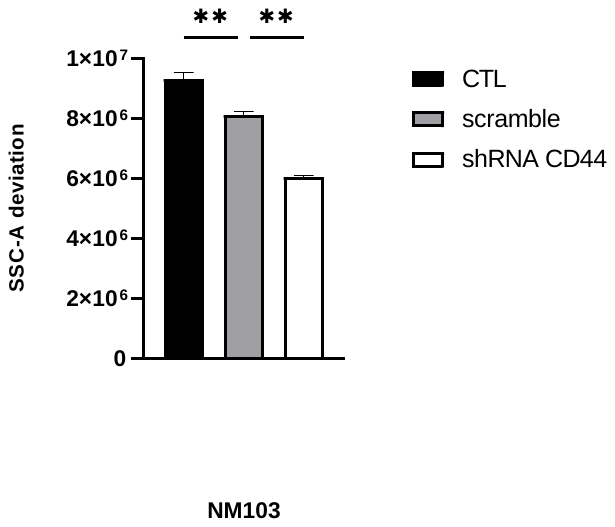

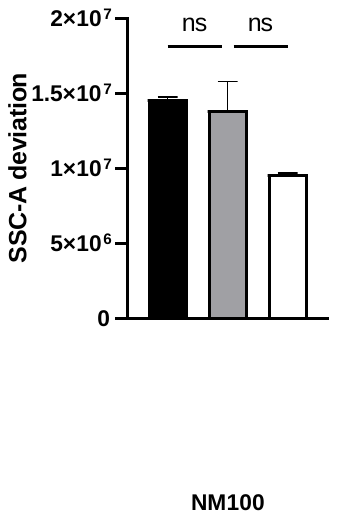

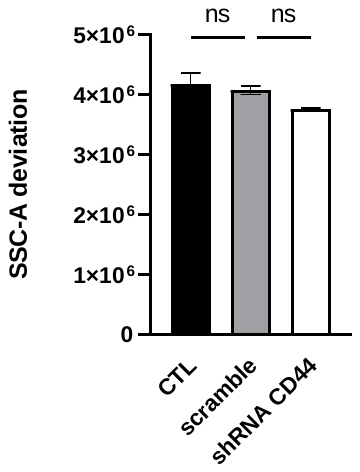

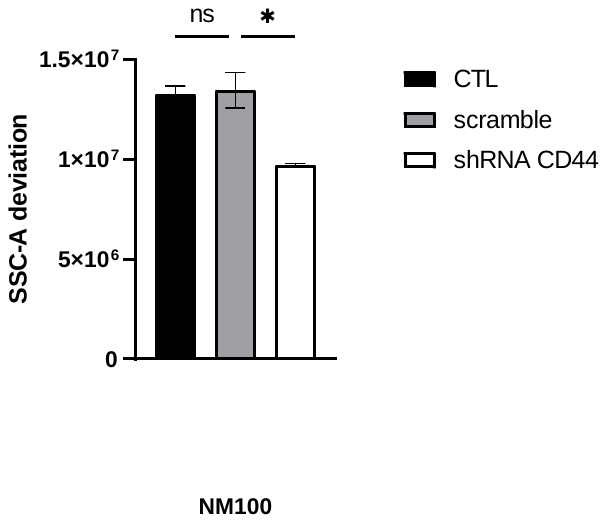

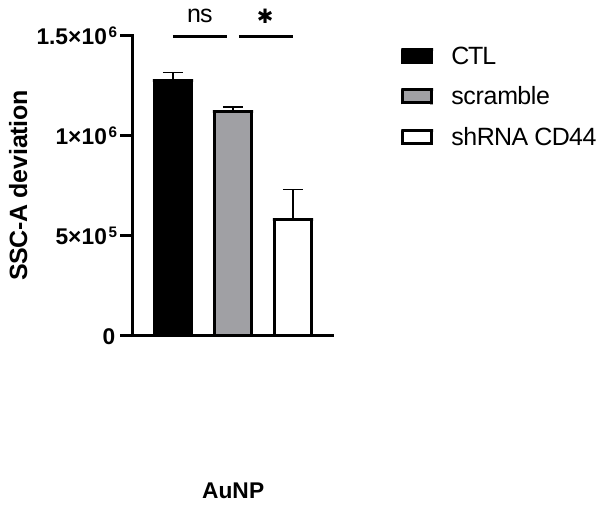

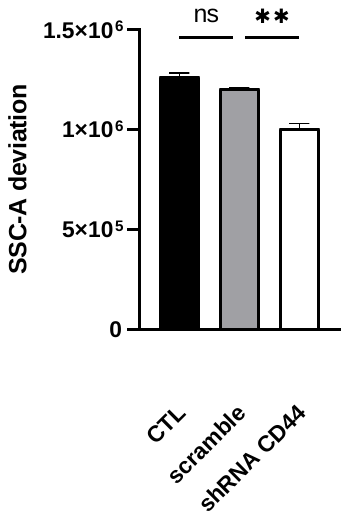

C

Replication 1

Replication 2

Replication 3

Replication 1

Replication 2

Replication 3

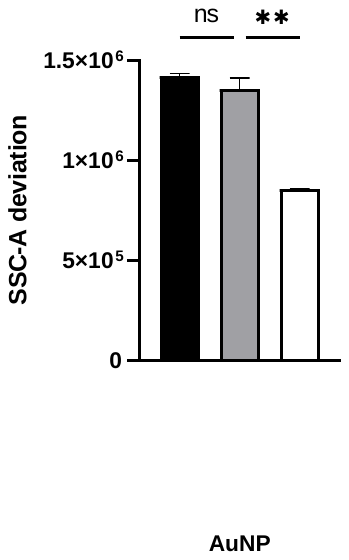

B

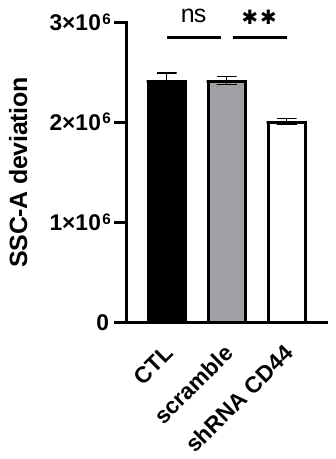

Replication 1

Replication 2

Replication 3

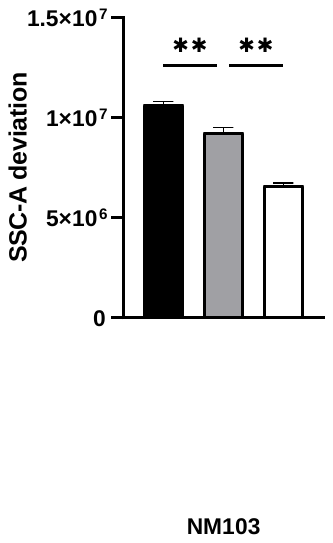

**Supplementary Figure 4 :** Replicate experiments showing how CD44 expression modification affects NP uptake

**A.** SSC-A deviation signals from MDA-MB231 cells expressing scramble or anti-CD44 shRNAs exposed to NM100 TiO_2_ NPs. The cells were exposed for 24 h to a NP concentration of 16 µg/cm², and suspensions were subjected to flow cytometry analysis. The graph represents the mean of data from 10000 cells, in duplicates. Statistical significance is validated by Ordinary One-way ANOVA and Sidak correction. Ns=non significant, *indicates a p-value < 0.05. **B.** SSC-A deviation signals from MDA-MB231 cells expressing scramble or anti-CD44 shRNAs exposed to NM103 TiO_2_ NPs. The cells were exposed for 24 h to a NP concentration of 16 µg/cm², and suspensions were subjected to flow cytometry analysis. The graph represents the mean of data from 10000 cells, in duplicates. Statistical significance is validated by Ordinary One-way ANOVA and Sidak correction. Ns=non significant, ** p-value<0.01. **C.** SSC-A deviation signals from MDA-MB231 cells expressing scramble or anti-CD44 shRNAs exposed to gold NPs. The cells were exposed for 24 h to a NP concentration of 8 µg/cm², and suspensions were subjected to flow cytometry analysis. The graph represents the mean of data from 10000 cells, in duplicates. Statistical significance is validated by Ordinary One-way ANOVA and Sidak correction. Ns=non significant, *indicates a p-value<0.05, ** p-value<0.01.

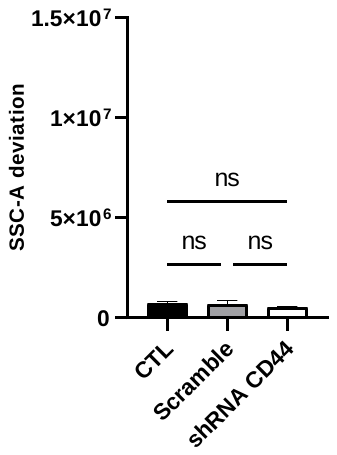

NM100

NM103

No NP

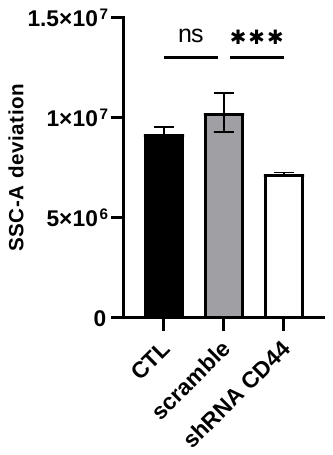

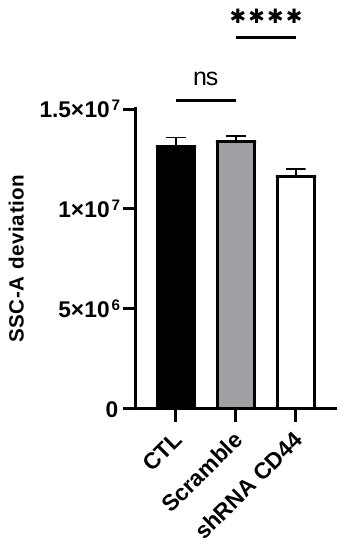

**Supplementary Figure 5 :** CD44 expression modification affects NP uptake in MCF10A-M line.

SSC-A deviation signals from MCF10A-M cells expressing scramble or anti-CD44 shRNAs, unexposed to NPs (left panel), exposed to NM100 TiO_2_ NPs (middle panel) or to NM103 TiO_2_ NPs (right panel). The cells were exposed for 24 h to a NP concentration of 16 µg/cm², and suspensions were subjected to flow cytometry analysis. The graph represents the mean of data from 10000 cells, in two biological replications of duplicates. Statistical significance is validated by Ordinary One-way ANOVA and Tukey’s correction. Ns=non significant, ***indicates a p-value<0.0005, ****indicates a p-value<0.0001.

.

NM100

NM103

AuNP

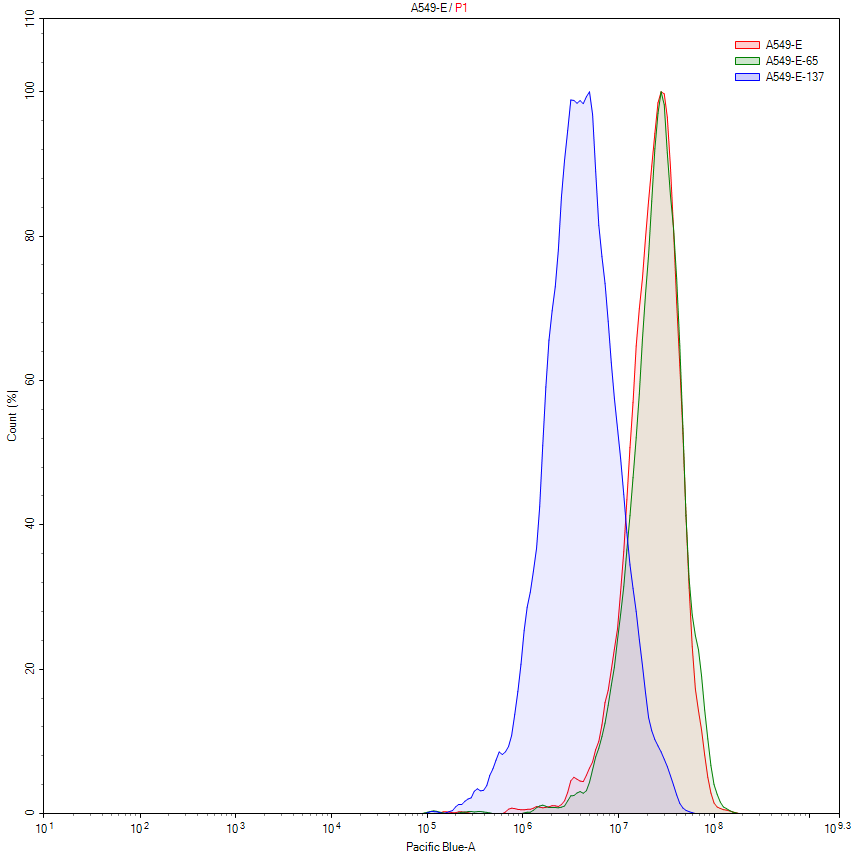

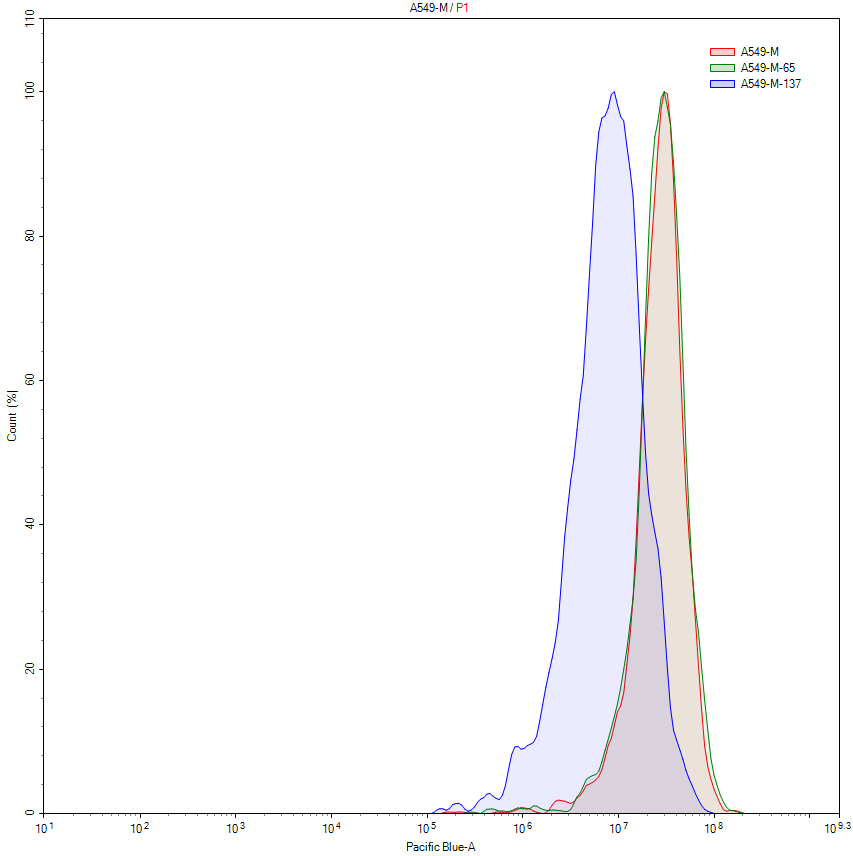

scramble

shRNA CD44

CTL

scramble

shRNA CD44

CTL

Count

CD44

Count

CD44

A549

A549-M

A

B

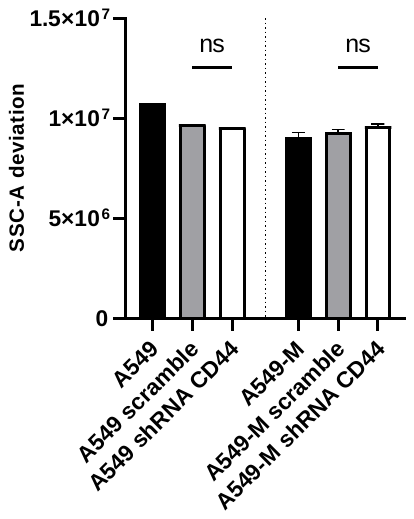

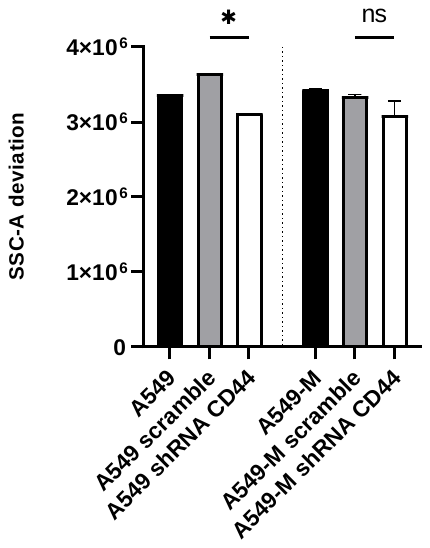

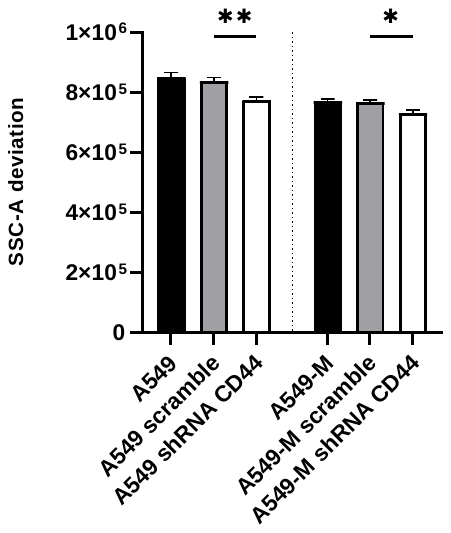

**Supplementary Figure 6 :** CD44 expression modification affects NP uptake in A549 and A549-M line.

**A.** Flow cytometry phenotyping of A549 (left) and A549-M (right) cells after expression of shRNAs targeting all isoforms of CD44. Cells were transduced with lentiviral particles encoding scramble or anti-CD44 shRNAs and labelled with CD44-BV421 antibodies. **B.** SSC-A deviation signals from A549 and A549-M cells expressing scramble or anti-CD44 shRNAs, exposed to NM100 TiO_2_ NPs (left panel), to NM103 TiO_2_ NPs (middle panel) or to gold NPs (right panel). The cells were exposed for 24 h to a NP concentration of 16 µg/cm² (TiO_2_ NPs) or 8 µg/cm^2^, and suspensions were subjected to flow cytometry analysis. The graph represents the mean of data from 10000 cells, in duplicates. Statistical significance is validated by Ordinary One-way ANOVA and Sidak’s test. Ns=non significant, *indicates a p-value<0.05, **indicates a p-value<0.01.

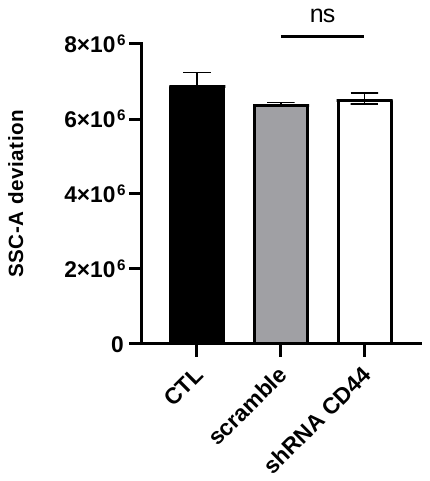

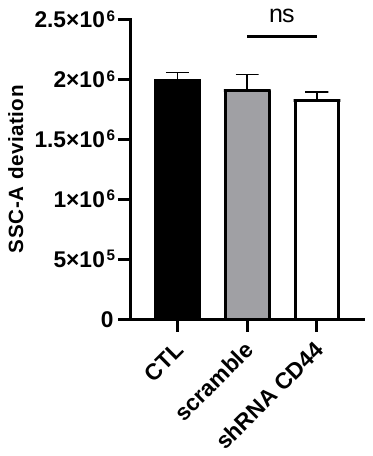

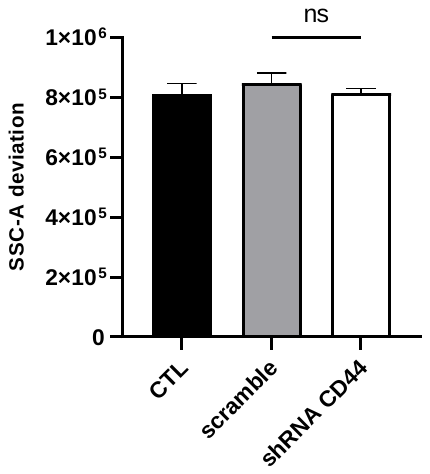

NM100

NM103

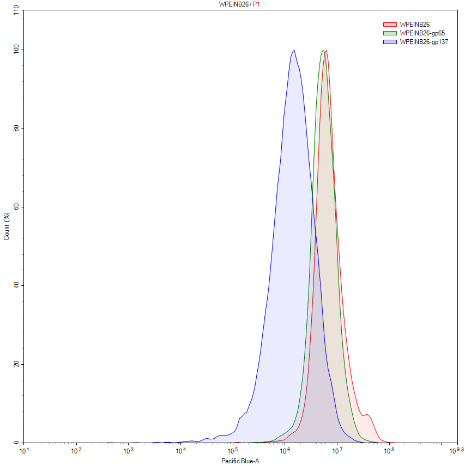

AuNP

scramble

shRNA CD44

CTL

WPE1-NB26

Count

CD44

A

B

**Supplementary Figure 7 :** CD44 expression modification affects NP uptake in WPE1-NB26 line.

**A.** Flow cytometry phenotyping of WPE1-NB26 cell line after expression of shRNAs targeting all isoforms of CD44. Cells were transduced with lentiviral particles encoding scramble or anti-CD44 shRNAs and labelled with CD44-BV421 antibodies. **B.** SSC-A deviation signals from WPE1-NB26 cells expressing scramble or anti-CD44 shRNAs, exposed to NM100 TiO_2_ NPs (left panel), to NM103 TiO_2_ NPs (middle panel) or to gold NPs (right panel). The cells were exposed for 24 h to a NP concentration of 16 µg/cm² (TiO2 NPs) or 8 µg/cm^2^, and suspensions were subjected to flow cytometry analysis. The graph represents the mean of data from 10000 cells, in duplicates. Statistical significance is validated by Ordinary One-way ANOVA and Sidak’s test. Ns=non significant.

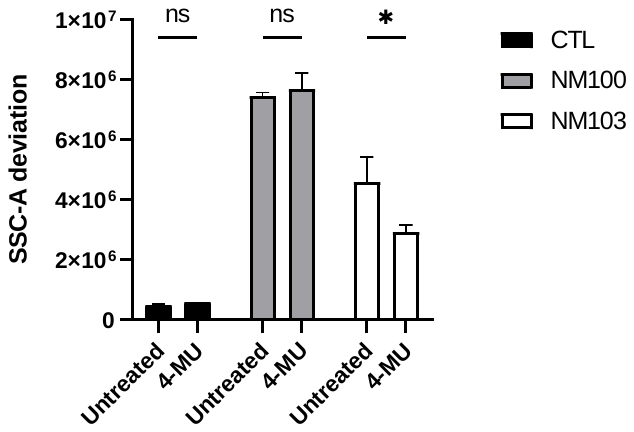

MDA-MB231 cells

A

T47D cells

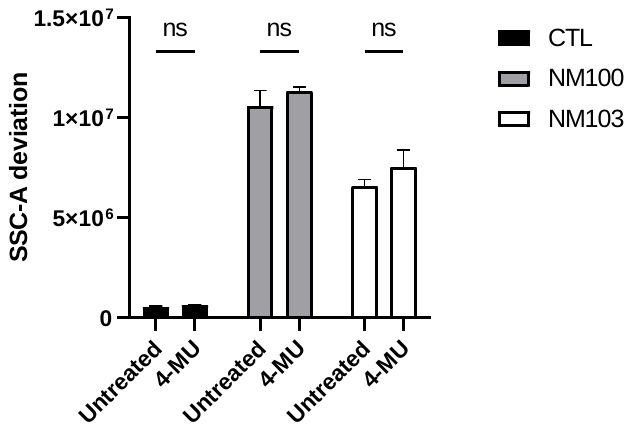

HMLE-E cells

HMLE-M cells

B

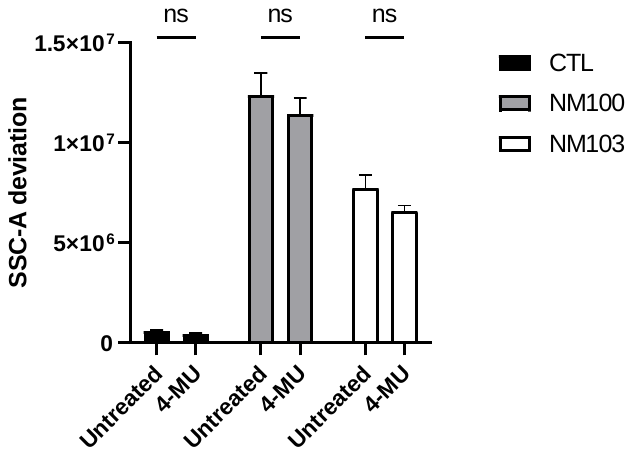

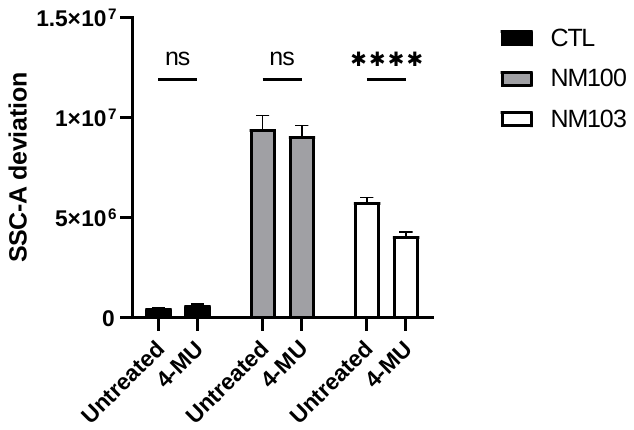

**Supplementary Figure 8 :** Effect of 4-methylumbelliferone on nanoparticles uptake in T47D and MDA-MB231 cells.

**A.** SSC-A deviation signals from T47D and MDA-MB231 cells exposed to NM100 or NM103 TiO_2_ NPs, after treatment with 4-methylumbelliferone. The cells were treated with 4-MU for 24 h and exposed for 24 h to a NP concentration of 16 µg/cm², and suspensions were subjected to flow cytometry analysis. The graph represents the mean of data from 10000 cells, in duplicates. Statistical significance is validated by two-way ANOVA and Sidak’s test. Ns=non significant, *p-value<0.05. **B.** SSC-A deviation signals from HMLE-E and HMLE-M cells exposed to NM100 or NM103 TiO_2_ NPs, after treatment with 4-methylumbelliferone. The cells were treated with 4-MU for 24 h and exposed for 24 h to a NP concentration of 16 µg/cm², and suspensions were subjected to flow cytometry analysis. The graph represents the mean of data from 10000 cells, in 4 replicates. Statistical significance is validated by two-way ANOVA and Sidak’s test. Ns=non significant, ****p-value<0.0001.
